# Unraveling the Palindromic and Non-Palindromic Motifs of Retroviral Integration Site Sequences by Statistical Mixture Models

**DOI:** 10.1101/2022.10.26.513837

**Authors:** Dalibor Miklík, Jiří Grim, Daniel Elleder, Jiří Hejnar

## Abstract

A weak palindromic nucleotide motif is the hallmark of retroviral integration site alignments. Previously, the motifs were explained by an overlap of the non-palindromic motif being present on one of the half-site of targeted sequences. Here, we applied multicomponent mixture models to integration site sequences of diverse retroviruses. We demonstrate that the weak palindromic motifs result from a combination of independent sub-motifs restricted to only a few positions proximal to the site of integration. The sub-motifs are formed by either palindrome-forming nucleotide preference or nucleotide exclusion. Using the mixture models, we also identified HIV-1-favored palindromic sequences in Alu repeats serving as hotspots for integration. Our work presents a novel statistical approach to the analysis of retroviral integration site sequences, which can form a valuable tool in the analysis of DNA motifs. The presented results shed new light on the selection of target site sequences for retroviral integration.

## Main

Retroviruses integrate the DNA copy of their genome into the genome of the host. The distribution of proviruses is genome-wide with global preferences toward specific chromatin states and genomic features ^1–7^. Globally, tethering to chromatin by cellular proteins drives the preferences ^8–13^. Locally, the occupancy of nucleosomes and the physical attributes of DNA affect the target site selection ^14–18^.

Retroviral enzyme integrase (IN) is the key protein that processes viral DNA ends and catalyzes strand transfer reaction (STR). During the STR, 3’ hydroxyl groups of viral DNA attack phosphodiester bonds of target DNA (tDNA) leading to a covalent connection of the host and viral DNA. Typically, the STR sites on both strands of tDNA are 4 to 6 nucleotides apart depending on the retrovirus. Structural studies demonstrated that IN assembles into symmetric multimeric complexes that bend the tDNA enabling the STR inside a widened major groove of tDNA ^18–27^.

After the alignment of preintegration tDNA sequences, a weak palindromic motif arises as a hallmark of sites recognized by IN ^28–33^. The appearance of the palindromic motifs is dependent on IN amino acids that are in contact with tDNA ^19,34–38^. Hence, the selection of the precise position for integration exhibits a pattern consistent with the IN-dictated preference for tDNA composition. Given the symmetry of IN multimers, the preference for palindromic motifs may be expected. In addition, the palindromic motif is observed at target sites of other transposable elements ^39–44^ and is a marker of binding sites of some transcription factors ^45^.

Palindromes are, however, rarely present in individual tDNA sequences. It was suggested that the weak palindromic motif results from the non-palindromic motif being present on either tDNA strand, i.e. on one half-site of the tDNA ^46^. According to this hypothesis, the statistical distribution mixture model of two reverse-complementary components was proposed to decompose the set of sequences. In this way, the constrained model necessarily splits the populational motif into a pair of reverse-complementary sub-motifs.

Here, we have used unconstrained mixture models of multiple (*>* 2) components to unravel heterogeneous sub-populations of sequences targeted by retroviral integration. We show that the integration site motifs are formed by an overlap of the motifs generated by either the preference for or exclusion of particular nucleotides. Moreover, using the multicomponent mixture models we identified palindromic sequence-defined hotspot of HIV-1 integration located in Alu repeats.

## Results

### EM algorithm uncovers palindromic and asymmetric sub-motifs in retroviral IS

To uncover sequence motifs at retroviral integration sites (IS), we estimated the mixture models of IS data sets utilizing the expectation-maximization (EM) algorithm. We collected data representing IS of human immunodeficiency virus type 1 (HIV) ^47^, human T-cell leukemia virus type 1 (HTLV) ^46^, maedi-visna virus (MVV) ^25^, Moloney murine leukemia virus (MLV) ^3^, prototype foamy virus (PFV) ^48^ and avian sarcoma and leukosis virus (ASLV) ^49^ (Extended Data Table 1).

The statistical properties of each IS set are characterized by a distribution mixture including a fixed number of components. A component is defined by i) nucleotide position probability matrix (PPM) and ii) by component weight representing the proportion of mixture covered by the component. To portray the PPMs we developed a sequence logo based on the Kullback–Leibler information divergence (KLID) that emphasizes the divergence of the observed frequencies to the genomic probabilities of nucleotides. Additionally, we calculated the palindromic defect (PDef) of PPMs. The PDef corresponds to the distance from self-reverse-complementarity and is equal to zero for palindromic PPMs. The PPMs representing entire retroviral IS sets (PPM0) exhibited low KLID values in sequence logo (Fig. 1A) and close-to-zero PDefs (Fig. 1B) confirming the weak palindromic motifs at IS alignments.

**Figure 1:**
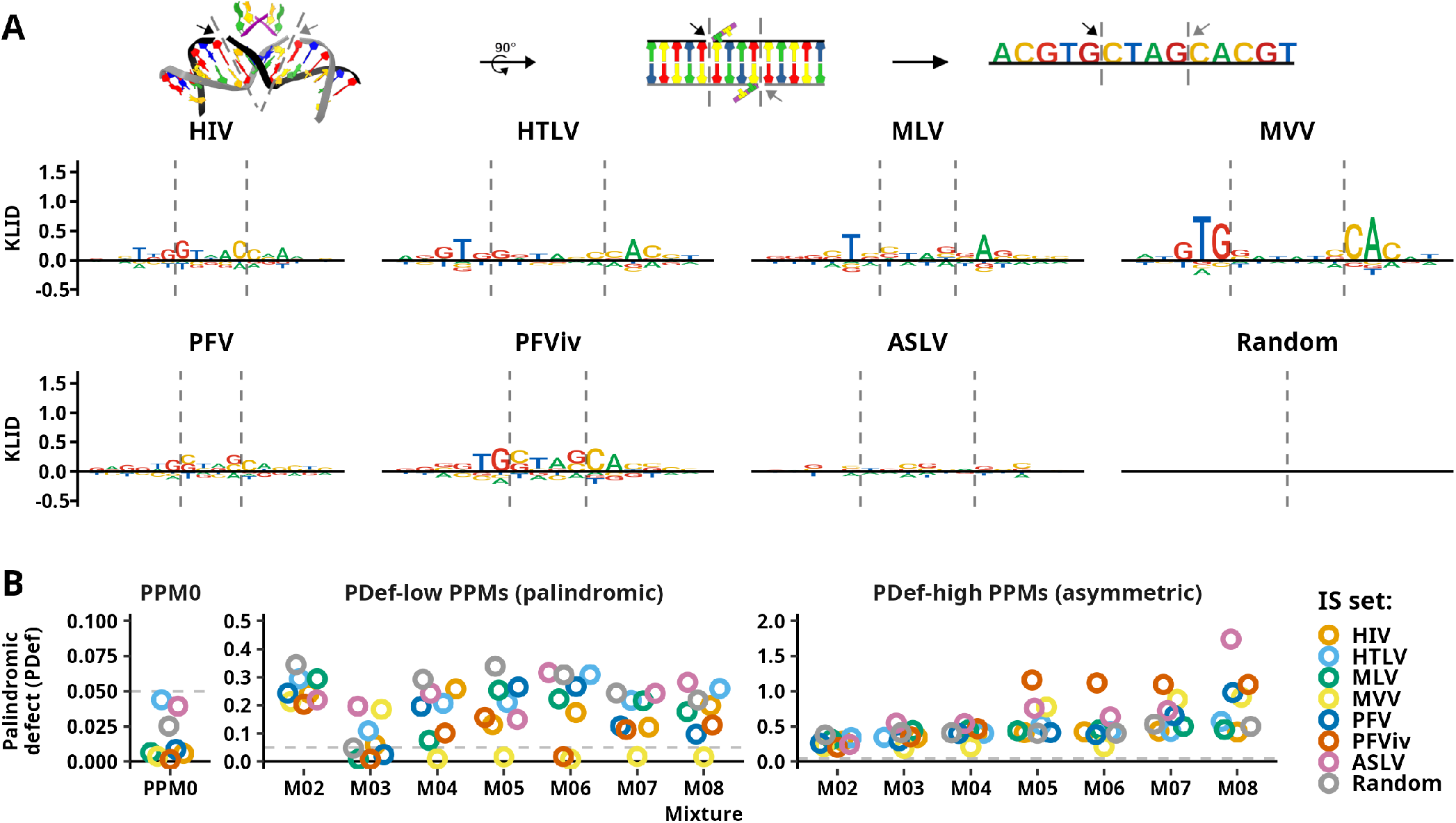
Identification of palindromic and asymmetric mixture components. A) Scheme of retroviral integration and sequence logos representing complete sets of retroviral IS and a set of random genomic sequences. The vertical dashed lines mark the sites of strand transfer reaction (STR), and the arrows point to the sites where the STR takes place. Logos represent IS sequences spanning 8 nucleotides to each side from the center of the sequences. B) Representation of palindromic defect (PDef) of selected component PPMs. Single PPM is shown for each retroviral IS component mixture. On the x-axis, mixtures are ordered by the number of components from two-component (M02) to eight-component (M08) mixture. PPM0 represents the PPMs of the complete IS set (left panel). The middle panel shows PPMs displaying the lowest PDef in the mixture. The right panel shows PPMs with the highest PDef in the mixture. For easier comparison between the panels, the horizontal dashed line representing PDef = 0.05 is shown.

The PDef was calculated for each PPM of estimated mixtures (Extended Data Fig. 1). While the two-component (M02) mixtures were absent of PDef-low PPMs (Fig. 1B), the PDef-low PPMs appeared in the mixtures containing three or more components. For instance, each of the M03 to M08 MVV mixtures contains a highly palindromic component. Asymmetric (PDef-high) PPMs are also present in M03 and other multicomponent mixtures.

The palindromic and asymmetric nature of the components was confirmed by the examination of the sequence logos (Extended Data Fig. 2B). The most palindromic components are often derived from M03 mixtures and (except for HIV) are enriched in A and T nucleotides with dominant T/A palindromic motif. In contrast, the most asymmetric PPMs are often derived from M07 or M08 mixtures and are enriched in G/C nucleotides (Extended Data Fig. 2C).

Initial analysis identified both asymmetric and palindromic components in estimated IS mixtures. While M02 mixtures preferably create pairs of asymmetric components, M03 mixtures seem to be selective toward the creation of at least one palindromic component. Both representations could still artificially reflect the overlap of sequences containing distinct motifs. The introduction of multicomponent mixtures is thus necessary to minimize the creation of artificial motifs.

### Multiple motifs are present in mixtures of IS sequences

To get the complete picture of the mixture composition, we examined the sequence logos of all components forming the mixtures (Supplementary Sequence Logos). Visual examination reveals the high heterogeneity of the sequence motifs both inside the mixtures and between the retroviral IS sets. Despite this heterogeneity, several motifs repeatedly appear among the components of independent mixtures. First, there is a clear separation of A-, T-, G-, or C-enriched components. The simple nucleotide content thus may be a common feature of some IS sequences. For instance, A/T-rich components are present in M08 mixtures of HIV (C04, C02), HTLV (C02, C05) and MVV (C03) IS sets (Fig. 2). The MVV M08 C03-like palindromic component appeared in every MVV IS mixture that included at least three components, covers 7-11% of the respective mixtures and lacks otherwise ubiquitous enrichment for palindrome-forming G/C nucleotides upstream to the STR sites. This observation suggests that A/T-rich sequences may form a special category of target substrates.

**Figure 2:**
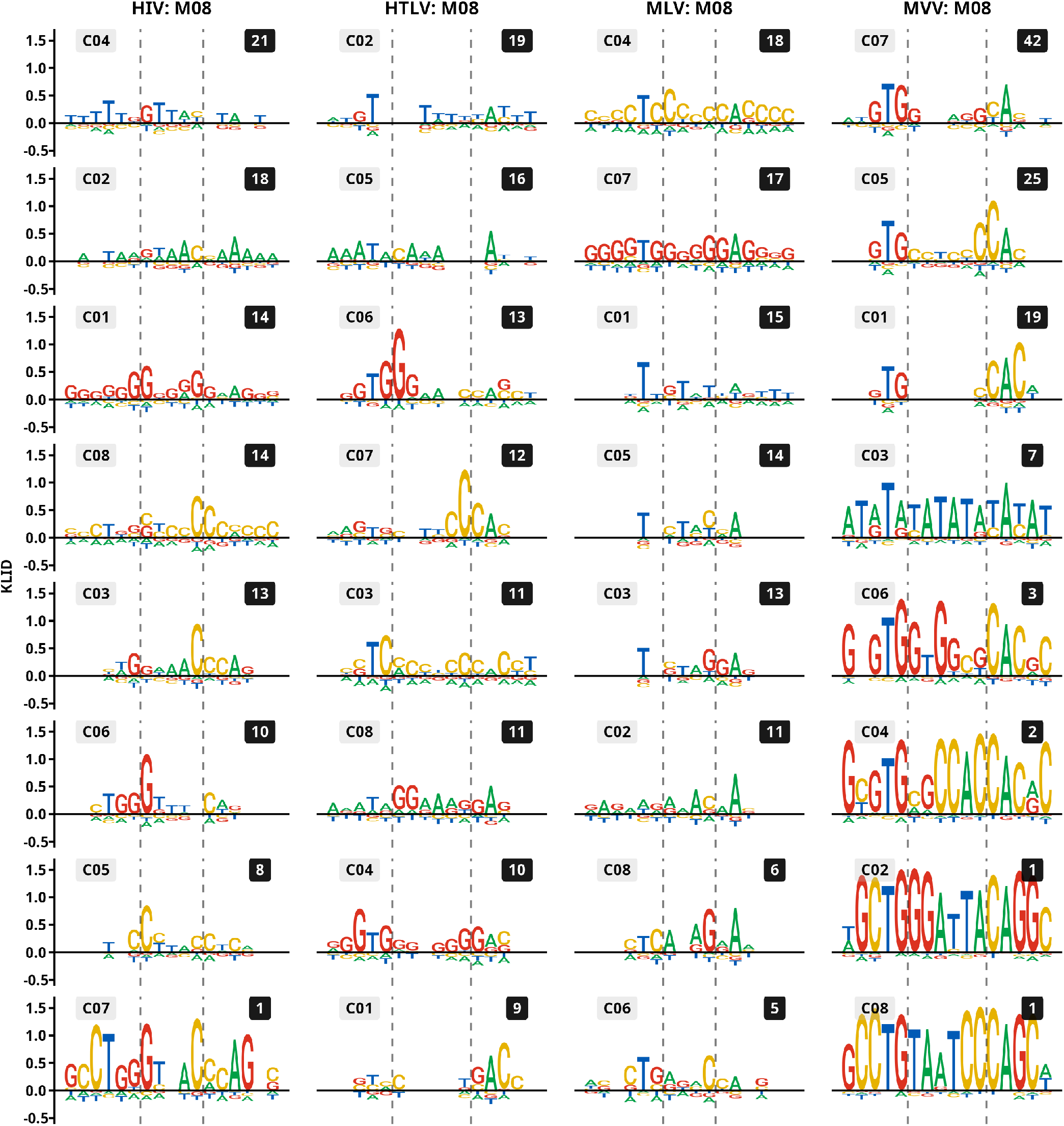
Sequence logos of complete component mixtures of retroviral IS sets. KLID sequence logos of the IS mixtures. Eight-component mixtures of HIV, HTLV, MLV, and MVV IS sets are shown as examples of highly decomposed mixtures formed by components displaying various motifs. Logos represent IS sequences spanning 8 nucleotides to each side from the center of the sequences. Vertical dashed lines mark sites where strand transfer reaction takes place.

The enrichment for certain nucleotides at specific positions was also repeatedly observed. The palindrome-forming enrichment of the T/A nucleotides at positions outside the inter-STR area is one frequently present motif. This palindromic motif is most prominent in MVV and MLV mixtures, where it is present even in G/C-rich components (Fig. 2, C04 and C07 of MLV and C07, C05 and C01 of MVV mixtures) and rarely segregates into an asymmetric motif. On the other hand, HIV and HTLV mixtures contain components with asymmetric enrichment of G/C nucleotides inside the inter-STR area (Fig. 2, HIV C03/C06 and HTLV C06/C07). The components form the bona fide asymmetric reverse-complementary pairs which together cover 20-40% of the respective mixtures. Other IS set-specific positional motifs like asymmetric G/C enrichment outside the inter-STR area (Fig. 2, HTLV C04/C01) or TA enrichment at central positions of the inter-STR area specific for *in vitro* IS (Supplementary Sequence Logos, PFViv M08 C02 and C06) were observed.

Examination of the whole mixture models revealed the complexity of the retroviral IS sets and validated the existence of both palindromic and asymmetric motifs in the multicomponent mixtures. The observation of motifs segregating into distinct components indicates the existence of independent separable sub-motifs. Importantly, the described motifs appear repeatedly in independently estimated mixtures adding to the reliability of the observed sub-motifs.

### Preferred nucleotide combinations at specific positions of IS sequences

To directly inspect the IS sequences for the motifs observed in estimated mixtures, we determined the frequencies of nucleotide combinations at complementary positions proximal to the STR sites (see Methods for detailed description; in text, the positions are labeled by square brackets).

First, we calculated KLID that indicates the enrichment for nucleotide combinations at each position (Fig. 3A, Extended Data Fig. 3). Generally, we observed that high KLID is caused by either a single dominant combination or, when the KLID is higher than a single contribution, by multiple nucleotide combinations. Upstream to the STR site, the palindrome-forming T-T combination is the dominant combination observed at positions [-2] or [-3] followed by the G-G combination. In contrast, at position [1], several combinations contribute to elevated KLID.

**Figure 3:**
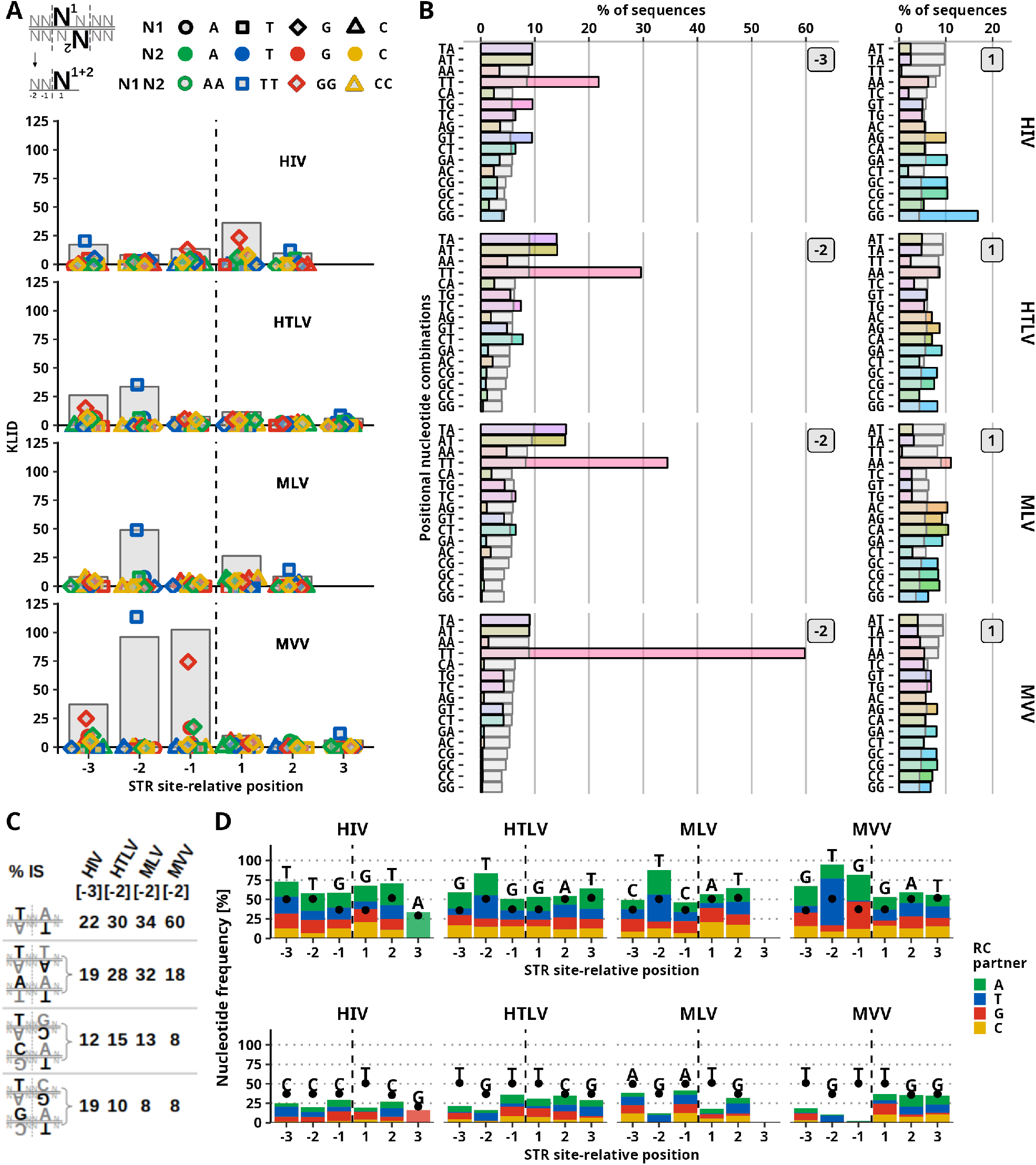
Preferred nucleotide combination at positions proximal to STR site. Frequency of nucleotide combinations at complementary positions of both DNA strands. A) KLID score of dinucleotide combinations at complementary sites marked by position relative to the STR site. The position of the STR site is marked by the vertical dashed line. Positions upstream to the STR site are marked with negative values. Gray bars represent the total KLID value at the position. Colored points represent individual contributions of each of the nucleotide combinations. Colors and shapes of points marking palindromic combinations are depicted in the legend. B) Frequency of sequences with marked positional nucleotide combination. Gray bars represent frequency observed in a control (random) set of sequences. The numbers right of the bars show position relative to the cleavage site. The left column represents the positions with maximum KLID score upstream of the cleavage site, and the right column represents the downstream-STR site-proximal position [1]. C) Representation of frequency of T-containing dinucleotide combinations at position [-3] of HIV and position [-2] of HTLV, MLV, and MVV IS sequences. D) Representation of frequencies of the most (top) and the least (bottom) frequent nucleotide at a given STR-site relative position. The colored area of the bar represents the frequency of nucleotide present at complementary STR site-relative position (i.e. position with the same number but on opposite DNA strand). Black points represent the nucleotide frequency in the random set of sequences.

The STR site-upstream T-T combination is present in 22% of HIV IS at position [-3] and in 27-60% of PFViv, HTLV, MLV, and MVV IS sequences at position [-2] (Fig. 3B, 3C, Extended Data Fig. 4). At the T-T enriched sites, frequencies of other nucleotides were increased when in combination with T. Strikingly, T nucleotide is observed at one of the sites forming [-2]/[-3] positions in up to 95% of IS sequences (Fig. 3D, top). On the other hand, G is the most disfavored nucleotide at the position (Fig. 3D, bottom). Besides the T-rich [-2] position, MVV IS sequences are enriched in G at position [-1] where G-G and G-A combinations are observed in 35% and 33% of IS sequences (Extended Data Fig. 4).

Position [1] is the site where asymmetric components of HIV and HTLV mixtures were enriched in G nucleotides. We observed G-G palindrome-forming combination at position [1] in 17% of HIV but only in 8% of HTLV IS sequences. More frequent are, however, the asymmetric G-A and G-C combinations that are each present in 20% of HIV and 18% and 16% of HTLV IS sequences. In total, 67% of HIV and 53% of HTLV IS sequences contain G at position [1]. The most striking, however, is the disfavor of combinations containing T nucleotide at position [1] observed also in MLV and PFV IS sequences that otherwise display only mild preferences for nucleotide combinations at the position (Fig. 3D bottom, Extended Data Fig. 5B).

In summary, we showed that at specific positions, the broken palindromes and T-exclusion form the globally weak motifs of retroviral IS. The broken palindromes appear upstream of the STR sites and are created by less preferred nucleotides disrupting the preferred palindrome-forming combinations. On the other hand, the exclusion of T at positions downstream to the STR sites causes the increase of other nucleotide frequencies at the position.

### Component mixture models identify hotspot of HIV-1 integration

Inspection of the multicomponent mixtures revealed low-weight components with strong motifs in the sequence logo. We hypothesize the components may describe small groups of highly similar sequences. To challenge the hypothesis, we characterized IS associated with the palindromic component C07 of HIV IS mixture M08 (Fig. 2).

We identified 1,587 C07-associated IS. The majority of the IS (96%) overlap with Alu repeats (Fig. 4A), which, when mapped to Alu consensus sequence ^50^, concentrated to two loci (Fig. 4B, Supplementary Table 1). Next, we reduced the IS sequences to the 13bp motifs formed by palindrome-forming positions [1], [-2], and [-3] (Fig. 2, 3A). Palindromic CT..G… C..AG motif was the most frequent motif in C07-associated sequences (Fig. 4C) as well as in intra-Alu IS sequences from the whole HIV IS set (Fig. 4E). Following our previous observations, most of the sequence motifs of the top ten intra-Alu-targeted motifs contained one half-site of the palindromic motif (5’-CT… G… −3’). While IS are enriched at the positions of the palindromic motif, enrichment is not observed in the proximity to the sites of the motif (Fig. 4F). These results suggest that the exact positions, not the surrounding loci, are locally preferred HIV integration hotspots.

**Figure 4:**
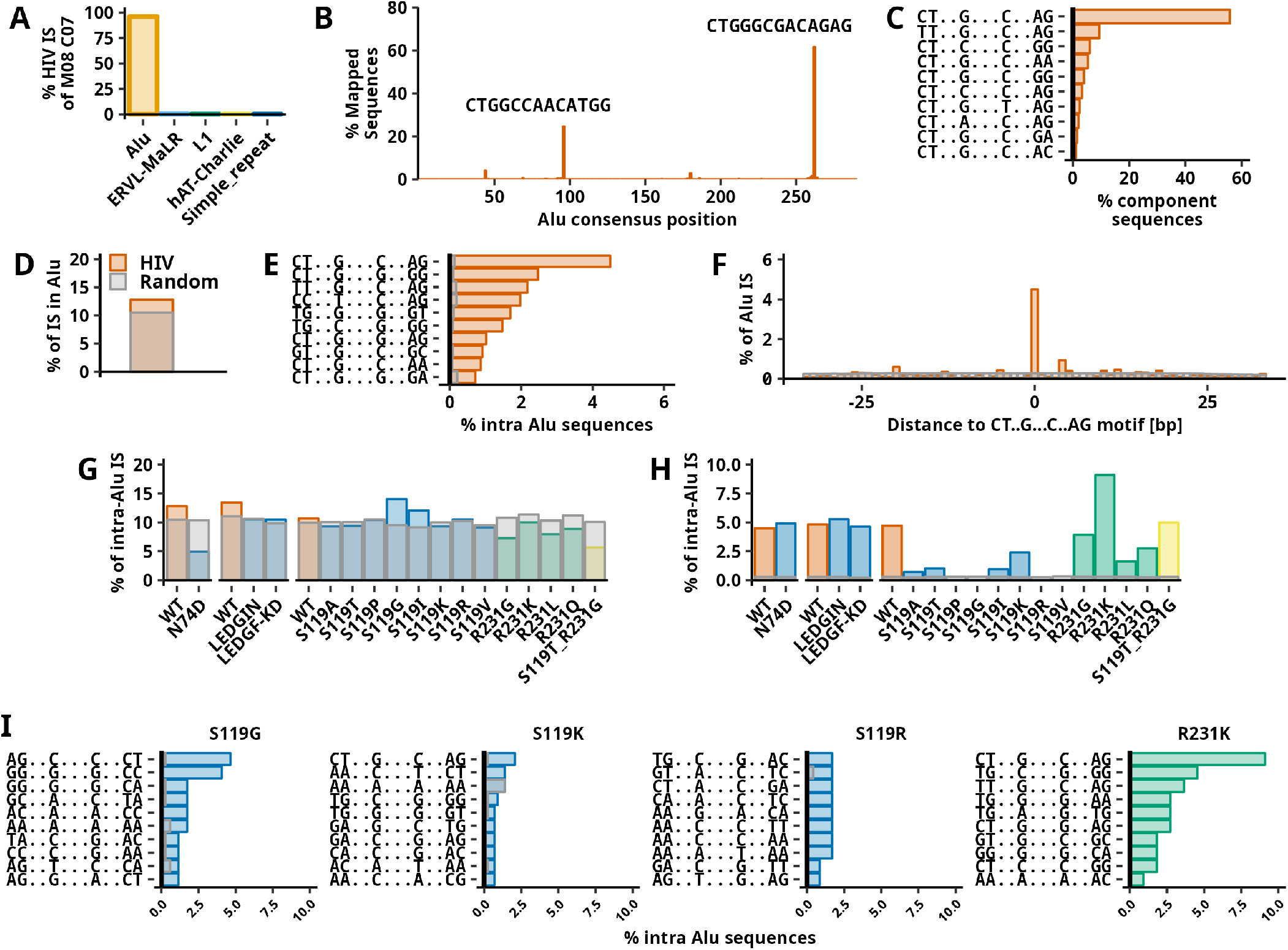
Characterization of HIV integration hotspot in Alu repeats. Panels A), B), and C) show the characterization of HIV-1 IS sequences associated with component C07 of a mixture M08. A) Barplot showing the frequency of 1,596 component-associated IS in repetitive elements. Five repeat families with the most IS are depicted. B) Intra-Al component-associated sequences mapped to Alu consensus sequence. The plot shows single-base binned consensus positions (length 290 bp) and a percentage of 1063 sequences mapped to individual positions of Alu consensus. Sequences of the two most frequent positions in the Alu consensus sequence are displayed. C) Frequency of sequence motifs in sequences associated with component C07. The ten most frequent sequence motifs are shown. In sequence motifs, dots (“.”) substitute any nucleotide. Panels D), E), and F) show data for HIV-1 obtained from the complete HIV IS set. D) Barplot showing a frequency of IS found in Alu repeats. The gray bar represents the expected random targeting of Alu repeats. E) Frequency of the sequence motifs among intra-Alu IS. Gray bars represent the mean expected frequency. The ten most frequent sequence motifs are shown. F) Frequency of integration into and in proximity to the most frequently targeted motif CT..G…C..AG. Plots show the frequency of intra-Alu IS 33 bp down- and upstream to the motif relative to Alu repeat orientation. Panels G), H) and I) show results of the same analysis as shown for panels D), E) and F) performed for different studies. Each experimental set contains a control with normal (WT) integration preference. N74D is HIV capsid mutation, LEDGIN marks the usage of LEGFF/p75-IN interaction inhibitor, LEDGF-KD transduction of cells where LEDGF/p75 protein is knocked-down. The right part of the bar plots represents IN mutants. Mutants of S119 are blue, mutants of R231 are green, and mutant of both S119 and R231 is yellow. G) Frequency of HIV IS in Alu repeats. H) Frequency of intra-Alu repeats in palindromic motif CT..G…C..AG. I) Frequency of ten most frequent sequence motifs.

To test the effect of preintegration complex tethering to chromatin, we extracted HIV IS data sets where capsid-CPSF6 (N74D) ^47^ or IN-LEDGF/p75 (LEDGIN treatment or LEDGF/p75 KD) ^51^ interactions were disrupted. Uniform preferential integration into the CT..G… C…AG motif was documented in each of the analyzed IS sets (Fig. 4G, H, Extended Data Fig. 6,7). The substitutions of IN amino acids ^34^ however impaired the targeting of the motif. It is noteworthy that every S119 mutant analyzed displayed the shift away from the palindromic hotspot caused by either more random selection for nucleotide combination at IS (S119G/I/K/R/V) or enhanced preference for nucleotide combinations at certain positions relative to IS (S119A/T) (Extended Data Fig. 8).

Using the information from the mixture components, we identified a local integration hotspot of HIV. The hotspot has a form of CT..G… C..AG palindromic motif which is preferentially located near the 3’ end of Alu repeats. The sequence hotspot is observed irrespective of the tethering of the HIV preintegration complex to chromatin and is an intrinsic feature of HIV IN. Under conditions not disrupting integration preference, about 1 in 175 HIV IS (Extended Data Table 2) is on average located at the hotspot.

## Discussion

The selection of the tDNA sequence is the last step in the complex process of retroviral integration site determination. Using multicomponent mixture models, we unraveled the sub-motifs present in the sequences targeted by retroviral integration. Results retrieved from our models showed that the weak palindromic motif of retroviral IS is formed by separate motifs formed by broken palindromes and nucleotide exclusion (Fig. 5). In addition, we demonstrated that the models can be used to identify locally preferred sequences of retroviral integration. In this way, we characterized HIV-1 palindromic hotspot in Alu repeats.

**Figure 5:**
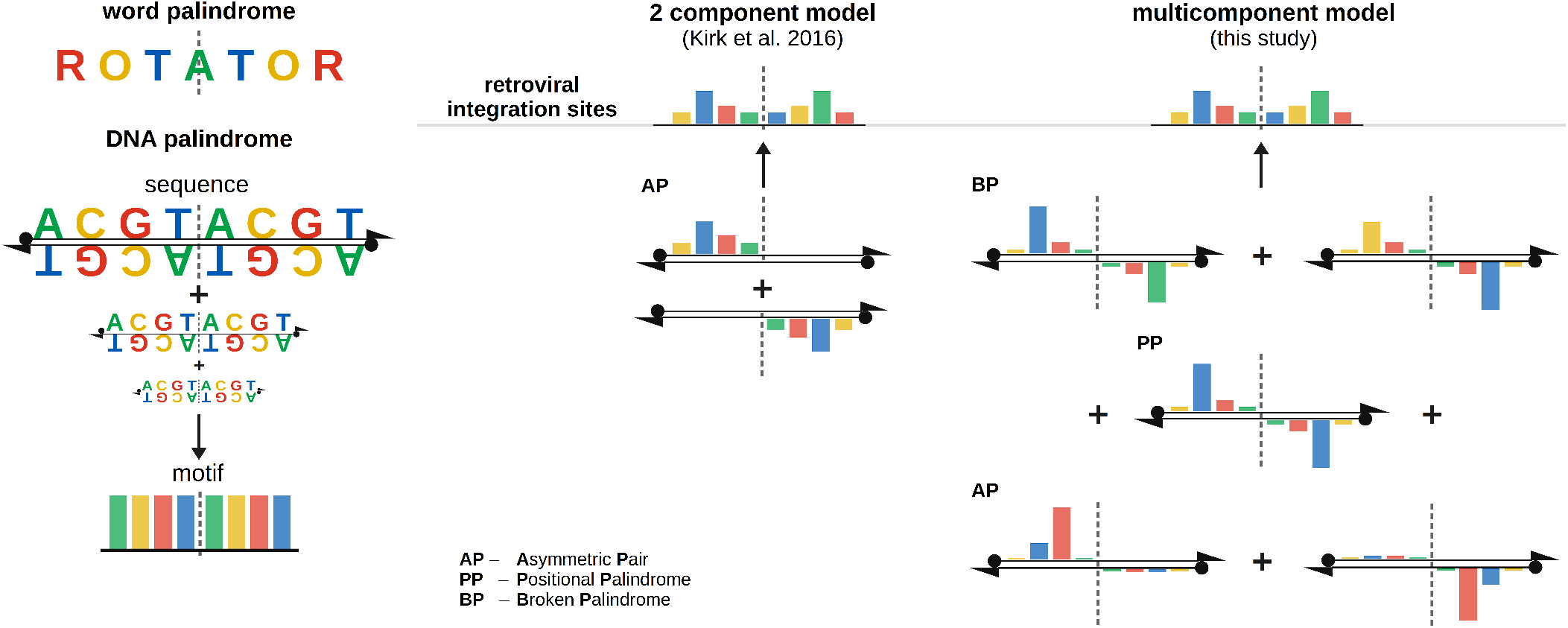
Summary of the models explaining weak palindromes at retroviral integration sites. DNA palindromes, like the word palindromes, are sequences that read the same forward and backward (from Greek “palin” again and “dramein” to run). A palindromic DNA motif composed of two reverse-complementary half-sites appears when palindromic sequences are aligned. The weakness of the palindromic motifs characteristic for integration sites of retroviruses suggests the low frequency of the palindrome in aligned DNA sequences. As the majority of the targeted DNA sequences are not palindromic, the phenomenon was previously explained by the model where an asymmetric motif appears on either of the half-site of the target DNA. Our model incorporates multiple independent motifs including preferred and often broken palindromic motifs as well as asymmetric motifs. Particular motifs appear at specific positions of target DNA and their frequency is specific for a given retrovirus.

A previous study suggested that the weak palindromic motif appears as a consequence of a non-palindromic motif occurring on one of the tDNA half-sites ^46^. First, the mixture model used was imposed to create two reverse-complementary motifs. Secondly, we demonstrated that even the unconstrained two-component mixture models are selective toward the creation of asymmetric pairs of motifs. The mixture models consisting of at least four components were needed to make convenient conclusions about the motifs in sets of retroviral IS. Although our approach may also be skewed toward the selection of mixtures containing reverse-complementary pairs of components (see Methods), we conclude that the asymmetric nature of IS sequences is better described by the existence of multiple independent motifs than by a single non-palindromic motif.

One explanation for the nucleotide preferences might reside in specific interactions at the IN-tDNA interface. About 3 bp upstream to the STR site, structurally conserved intrusion into the minor groove of small amino acids like serine (HIV, RSV), proline (HTLV, MVV, MLV ^36^, MMTV ^18^), and alanine (PFV) was documented. Albeit no nucleotide-specific interactions seem to be present at the position, mutations of HIV-1 IN S119 ^34,35,37^ affect the preference of IN target site selection. Interestingly, substitutions of S119 can either enhance or cancel the preference for nucleotides at positions proximal to the STR site and can disrupt the T-exclusion at STR-downstream position. The identity of the minor-groove intruding amino acid is thus the major determinant of IN preference for tDNA composition.

The exclusion of T at the position downstream to the STR site was previously explained by hindering the target phosphodiester by the C5-methyl group of T ^19^. Additional preference for G at the position of HIV-1 IS could potentially be explained by the interaction of G N5 with the major groove-invading arginine (R231) ^23^. Although R231G mutation was previously associated with mild global retargeting of HIV-1 integration ^34^, we did not observe altered targeting of Alu hotspot nor altered nucleotide preferences of IN R231 mutants. In addition, no significant preference for G downstream to the STR site is present in IS sets of PFV and MVV, where the major groove-invading arginine is present. Moreover, obstruction of the STR by the T C5-methyl group may be overcome by substitutions of other amino acids. Hence, the contribution of major-groove invading arginine to G preference at the position is disputable.

Since there are no clear correlates of IN-tDNA interface composition and the strength of tDNA motifs, the motif may reflect a need for tDNA to adopt specific conformation. Yet, there is no major distortion of B-form tDNA structure at the position occupied by preferred T ^18^. Also, the number of documented H-bond-forming interactions between IN and tDNA does not correlate with the preference for T-palindrome upstream to the STR.

Given the symmetry of retroviral intasome, the existence of non-palindromic sub-motifs is an interesting feature of the tDNA sequences. Imperfect palindromes are known to regulate the binding of transcription factors where changes in one half-site affect the binding of the transcription factor to the other half-site of the motif ^45^. Unlike the motifs of transcription factors, retroviral IS motifs are limited to a few positions and are probably not the consequence of the sequence-specific IN-tDNA interactions. However, we showed that the presence of the preferred nucleotide at one half-site of tDNA can increase the chances of the less preferred nucleotide appearing at the other half-site of tDNA. We thus consider the model where triggering the STR after tDNA recognition by IN can act asymmetrically.

*In vitro* experiments suggested that HIV-1 IN preference toward tDNA patterns shapes the local distribution of HIV-1 integration ^32,53^. Using the mixture models, we identified a group of highly similar sequences within *in cellulo* IS set. This hotspot was located to Alu repeats and represents the sequence with a palindromic pattern that might be an ideal template for HIV-1 integration ^32^. We observed the hotspot with constant frequency (1 in 175 IS) in independent IS sets and we demonstrated that the targeting of the hotspot is an attribute of HIV-1 IN. The existence of such a hotspot highlights the fact that, at least for HIV-1 integration, the sequence composition shapes the local distribution of proviruses and that scanning of tDNA by IN may take place before integration.

In summary, we demonstrated a novel statistical approach for the analysis of retroviral IS sequences which can form a valuable tool in the analysis of DNA motifs. So far, studies exploring tDNA composition approached the tDNA as the perfect palindrome or worked on the averaged global alignment of IS sequences. We propose that the demonstrated variability of target site sequences should be considered to understand the natural process of retroviral integration.

## Methods

### Integration Site Sequences

Sources of IS data are described in Extended Data Table 1. Only IS of HTLV ^46^ were obtained as genomic pre-integration sequences from supplementary data of the publication. For the rest of IS sets, genomic coordinates of LTR-proximal nucleotides were obtained from published coordinates, sets of mapped reads or raw sequencing reads. Details on the data processing can be found in Supplementary Methods. Briefly, genomic coordinates of nucleotides proximal to proviral LTRs were first obtained from available data. In the case of HIV data from Zhyvoloup et al. ^48^, IS coordinates were obtained from a publication supplementary table. Replicates of DMSO treated samples of wt and capsid N74D mutant were joined to create wt and N74D IS sets. Genomic coordinates of HIV IS data from Vansant et al. ^52^ were obtained from the supplementary table of Gene Expression Omnibus entry. wt, LEDGIN, and LEDGF-KD data sets were created joining bulk, GFP+ and GFP-samples from the identical treatment group. HIV IS coordinates from Demeulemeester et al. ^34^ were extracted from tables provided by the study’s authors. For each variant, data from the transduction of different cell lines were mixed. Coordinates of MVV IS from HEK293T cells ^25^ and PFV IS sets ^48^ were obtained from the Gene Expression Omnibus database. MLV ^3^ set was obtained from raw sequencing reads that were mapped to hg38 human genome assembly using bowtie2 ^54^. Only reads mapping from the first nucleotide and mapping uniquely to the genome were accepted. Data from CEF cells transduced by RCASC ^49^ were used as data representing ASLV integration. Mapped reads in SAM format were obtained from the authors of the study. Alignments were filtered and converted to BED format using SAMtools view ^55^ and BEDtools ^56^ bamtobed tools. Another ASLV IS set ^57^ was retrieved as raw reads and processed as IS raw reads of MLV and PFV. However, this IS set was not used in the main study as some uncertainties were observed during the analysis of mixture models (Supplementary Fig. 1).

Next, single-bp coordinates of LTR-proximal nucleotides in BED format were transformed to ranges spanning 13 bp to each side. As a result, ranges of length 26 (HTLV, MLV, MVV, PFV, and ASLV) and 27 nucleotides (HIV) were created. Sequences from respective genomes were obtained using BEDtools getfasta tools. FASTA files were then transformed into tables of sequences. For the purpose of mixture modeling, nucleotides were encoded to numeric strings where A = 1, C = 2, G = 3, T = 4, N = 5.

### Random genomic sequences and shuffled controls

Sets of randomly selected genomic sequences were created to estimate the expected frequencies of nucleotides at loci targeted by retroviral integration. First, 9,000 random genomic ranges of lengths 26 and 27 from hg19 and 10,000 ranges of length 26 bp from galGal4 were created using BEDtools random tool. Ranges were then transformed to FASTA sequences using the BEDtools getfasta tool. Sequences containing Ns were removed and sequences were then encoded to numeric strings as described for IS sequences. Sequences derived from the hg19 assembly were used as controls for all IS sets derived from the human genome including those mapped to the hg38 assembly. Background nucleotide frequencies used for the creation of the sequence logo were obtained as a mean nucleotide frequency across all positions of 26 bp random sequences-derived PPM obtained from hg19 genome assembly.

To control for the expected frequency of nucleotide combinations at complementary positions of the IS, the frequency of the combination was calculated in sets of random sequences derived from hg19 or galGal4 assemblies for human or chicken genomes, respectively.

To control for random targeting of genomic features, a BEDtools shuffle tool was used with a sample BED file and a file containing chromosome lengths of the targeted genome obtained from UCSC goldenpath (*https://hgdownload.soe.ucsc.edu/downloads.html*). In the case of shuffled controls created to IS inside genomic features (i.e. Alu repeats), -incl and -f 0.6 options were included in BEDtools shuffle command. BEDtools getfasta tool was used to retrieve genomic sequences of generated ranges. When stated, shuffling was repeated for x-times and target frequency was calculated as a mean of the frequency obtained in a particular iteration.

### The mixture of multiple product components

Given a collection of aligned nucleotide sequences of length N, the basic statistical description of the data follows from the relative frequencies of the four bases A,C,G,T at each position. We denote the set of bases ℬ = {*A, C, G, T*}. The probabilities of bases are usually presented in the form of a *position probability matrix* (PPM) of size 4 × N. Denoting N = {1, 2, …, *N*} we can write:

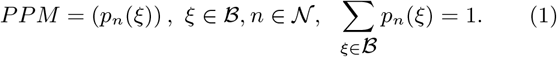

Here the rows of PPM correspond to the four bases A,C,G,T and the columns relate to the respective nucleotide positions n=1,2,…,N. In each column the probabilities sum to one. The *PPM* of the whole collection, in the following we use notation *PPM*_0_, can be viewed as a basic statistical model of IS sequences of the considered retrovirus.

In order to describe the statistical properties of IS sequence populations in a more specific way we have applied a general unconstrained mixture model of multiple components which can be estimated by means of EM algorithm. We have found that the reverse-complementary components tend to occur in mixtures spontaneously, without any enforced condition and the additional components may uncover other interesting properties of the IS sequences. For this purpose we assume the multivariate probability distribution *P* (***x***) in the form of a weighted sum of multiple product components *P* (*x* | *m*), *m* ∈ ℳ = {1, 2, …, *M*}:

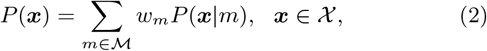

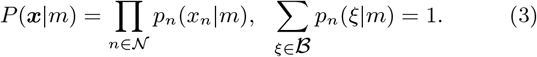

Here, *w*_*m*_ is the probabilistic weight of the *m*-th component and *p*_*n*_(*x*_*n*_ | *m*), *n* ∈ 𝒩 are the related component specific position probabilities of the bases A,C,G,T. In other words, the distributions *p*_*n*_(· | *m*), *n* ∈ 𝒩 define the component *PPM*_*m*_ of the underlying subpopulation.

### EM Algorithm

The distribution mixture (2) is a widely used general statistical model to approximate unknown discrete probability distributions. The standard way to estimate the mixture parameters *w*_*m*_, *P* (·|*m*), *m* ∈ ℳ is to maximize the log-likelihood criterion

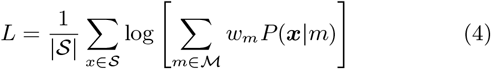

by means of the iterative EM algorithm ^52^.

Let 𝒮 be a collection of IS sequences of a retrovirus

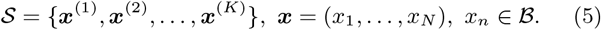

To compute the m.-l. estimates of the unknown parameters *w*_*m*_, *p*_*n*_(·|*m*) we repeat the basic EM iteration equations:

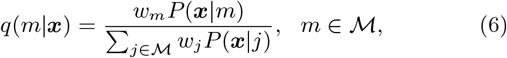

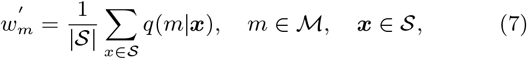

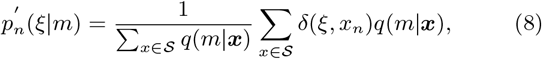

where | 𝒮 | is the number of sequences, 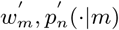 are the new parameter values and *δ*(*ξ, x*_*n*_) denotes the usual delta-function, i.e. *δ*(*ξ, x*_*n*_) = 1 for *ξ* = *x*_*n*_ and otherwise *δ*(*ξ, x*_*n*_) = 0.

The EM algorithm generates a nondecreasing sequence 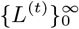 and the iterations are stopped when the relative increment of the criterion is less than a chosen threshold. The sequence 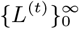 converges to a possibly local maximum which may be starting-point dependent. In order to decrease the risk of locally optimal solutions, the initial parameters *p*_*n*_(*ξ*|*m*) are usually generated randomly and the EM optimization is repeated several times with different initial values.

A difficult problem is a proper choice of the number of components *M*. It is highly data dependent and there is no reliable and easily applicable method to solve it. For this reason we have tested in our experiments different mixtures (*M* = 2 ÷ 12) with the aim to find the most probable model and to verify various aspects of mixture complexity.

### Extraction of component-associated sequences

Given the estimated distribution mixture (2) we can characterize any sequence ***x*** ∈ 𝒮 in terms of the conditional probabilities *q*(*m*|***x***). The conditional posterior weight *q*(*m*|***x***) can be interpreted as a measure of affinity of the sequence ***x*** to the m-th component or, in other words, as a membership value of ***x*** with respect to the *m*-th subpopulation. In this sense we can decompose the original data 𝒮 into sub-collections 𝒮_*m*_ according to the maximum values *q*(*m*|***x***), i.e. by means of Bayes formula. The membership value was calculated by custom R script and sequences with *q*(*m*|***x***) ≥ 0.9 were identified and separated from IS data set as a component-associated IS. In view of Eq. (7) the component weight *w*_*m*_ can be interpreted as an estimate of the relative size of 𝒮_*m*_

### Reverse-complement distance and palindromic defect

For any two position probability matrices *P*_1_, *P*_2_ the *reverse complement-distance* is defined as the sum of absolute differences of reverse-complementary position probabilities:

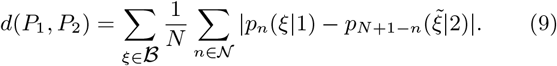

If the distance (9) is zero then, by definition, the two matrices *P*_1_, *P*_2_ are reverse-complementary.

If we apply the above formula (9) to one and the same matrix *P*_1_ = *P*_2_ = *P*_0_, then the corresponding value *d*(*P*_0_, *P*_0_) can be viewed as a measure of violation of the palindromic condition, i.e. as a *palindromic defect* of the matrix *P*_0_. We denote:

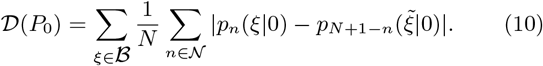

Here we ignore the central position if the number of positions N is not even. The palindromic defect 𝒟 (*P*_0_), (0 ≤ 𝒟 ≤ 2) is zero for any palindromic *P*_0_ and positive otherwise.

### Kullback-Leibler information divergenc

If *p*_*n*_(*ξ*|*m*), *n* ∈ 𝒩, *ξ* ∈ ℬ are the position probabilities of the m-th component and *p*_*n*_(*ξ*|0) the position probabilities of the background, then the Kullback-Leibler information divergence (KLID) of the two distributions *p*_*n*_(·|*m*), *p*_*n*_(·|0) at the position *n* ∈ 𝒩 is given by the formula

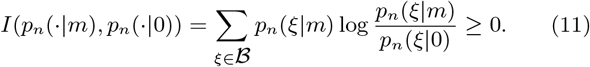

The KLID is nonnegative, equals zero only if the two distributions are identical and can be interpreted as a measure of information that is lost, if the IS probabilities *p*_*n*_(*ξ*|*m*) are replaced by the global background probabilities *p*_*n*_(*ξ*|0).

### Sequence logo based on KLID

A popular way to illustrate the related viral preferences is the well-known sequence logo derived from *PPM*_0_, which highlights the overall integration site motif. It is a histogram of N columns, each column displays proportionally the four probabilities of bases in descending order and the total height of the column is proportional to the Shannon information contained. The Shannon information is zero in the case of uniform probabilities and it is higher if some bases are preferred.

However, in the present context, the KLID formula is a more informative tool to illustrate the properties of mixture components graphically by a histogram and can be used as a sequence logo. Unlike the standard sequence logo, the value of the expression *I*(*p*_*n*_(·|*m*), *p*_*n*_(·|0)) can be viewed as a measure of information importance of the n-th position in the m-th component, in relation to the statistical properties of the background. The KLID formula (11) includes four terms that reflect the role of the four possible nucleotides. The term is positive if the IS probability *p*_*n*_(*ξ*|*m*) is greater than the global (background) probability *p*_*n*_(*ξ*|0) and it is negative, if it is smaller. In case of non-specific background distribution, characterized by *φ* _*mn*_ = 0 (see below), we have *p*_*n*_(*ξ*|*m*) = *p*_*n*_(*ξ*|0) and the related KLID value is zero. At each position of the histogram, the column height corresponds to the informativity *I*(*p*_*n*_(·|*m*), *p*_*n*_(·|0)). The four color parts are proportional to the respective contributions of the four nucleotides to the value of KLID. In contrast to the standard sequence logo, the contributions of the less probable nucleotides are displayed as negative, to emphasize their relation to the background.

The graphical representation of sequence logo based on KLID was created with the ggseqlogo R package ^58^. The functions of the package were modified to display the contribution of each nucleotide to KLID at the position. Given the set background probabilities (see chapter Structural mixture model), the maximum values for each nucleotide are 1.2226 for A, 1.5865 for C, 1.5714 for G, and 1.2271 for T.

### Structural mixture model

Obviously, the statistical analysis of a collection of aligned IS sequences based on a mixture model is a suitable approach to identifying the local integration site preferences of a retrovirus. However, in this way, the statitical properties of the near IS neighborhood may be influenced by random properties of the more distant parts of genomic sequences. In order to suppress the possible noisy influence of less informative positions of IS sequences we have used a structural modification of the product mixture (2). In particular, utilizing binary structural parameters *φ* _*mn*_ ∈ {0, 1} we can confine the estimation of component parameters only to some informative variables ^47^. If we define

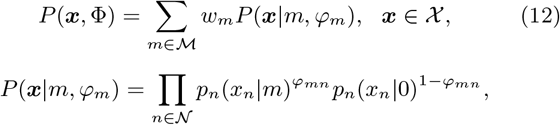

then, by setting the structural parameter *φ* _*mn*_ = 0, we can replace any component-specific distribution *p*_*n*_(·|*m*) by a common fixed “background” distribution *p*_*n*_(·|0). In our case we use the four global genomic probabilities *p*(*A*|0) = 0.2968, *p*(*C*|0) = 0.2049, *p*(*G*|0) = 0.2028, *p*(*T* |0) = 0.2955 as a background distribution:

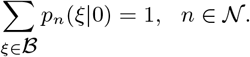

### Structural EM Algorithm

The structural mixture (12) can be optimized by the EM algorithm in full generality. In particular, making the substitution

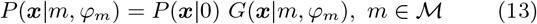

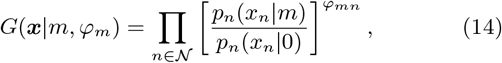

in (2), we can write the structural mixture in the form

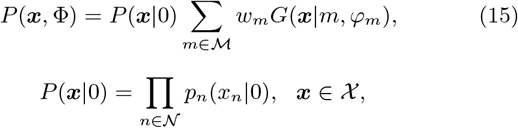

where *P* (***x***|0) is a fixed nonzero “background” probability distribution, and the component functions *G*(***x***|*m, φ* _*m*_) include binary structural parameters *φ* _*mn*_ ∈ {0, 1}. Obviously, by considering Eq. (2), the structural mixture model (15) can be viewed formally as a product mixture again.

By means of EM algorithm we can estimate both the component specific distributions *p*_*n*_(*x*_*n*_ | *m*) and the binary structural parameters *φ* _*mn*_, ^47^. For this purpose, we use the standard EM iteration equations (6) - (8). By means of substitution (15), we can write the structural log-likelihood criterion in the form

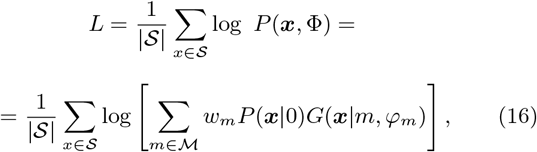

and, by using the formula (13), we can reduce the iterative Eq. (6) to informative variables only:

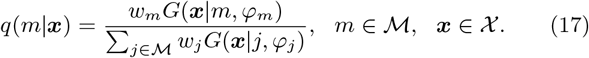

The next two Eqs. (7), (8) are unchanged. The only difference is that, in addition, we choose in each iteration the most informative parameters by means of the well known Kullback-Leibler information divergence.

In each iteration the KLID formula (11) is evaluated for all components and variables and the following weighted mean is used to derive a suitable threshold value:

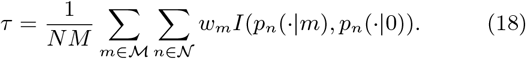

In the next iteration step only the sufficiently informative component distributions *p*_*n*_(·|*m*) are used satisfying the inequality:

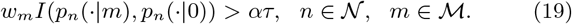

Here *α* is an optional coefficient which can be used to control the level of suppressed noise. By setting *α* = 0.0 we would estimate all component specific parameters, as in general mixture. The value *α* = 1.0 would eliminate all position distributions *p*_*n*_(·|*m*) of below-average informativity. In our case we have used *α* = 0.2 in order to replace only the low informative position distributions by background.

KLID is widely used in statistics and information theory as a measure of discrimination information. However, it should be emphasized that the application of KLID in the present context is not intuitively motivated, but directly follows from the monotonic condition of EM algorithm ^52^.

The EM iterations have been stopped when the relative increment of the criterion was less than 10^−5^. Again, in order to decrease the risk of locally optimal solutions, the EM algorithm has been initialized randomly and repeated 100-times with different initial values. Initially, the maximum of the log-likelihood has been used as a criterion, but in the final experiments the reverse-complement distances of all components were computed and the mixture containing pair of components with the lowest reverse-complement distance was selected. The minimum reverse-complement distance as criterion yields better interpretable results with comparable values of the log-likelihood criterion.

### Positional nucleotide combinations

To provide information about the frequency of nucleotide combinations at strand transfer reaction (STR) site-relative positions, STR site-relative PPMs were created, denoted as *PPM*_*STR*_. First, the alignments of IS sequences were split into equally long left *A*_*L*_ and right *A*_*R*_ half. The middle position (marked by 0 in PPM) is ignored if the original number of positions N is odd. The length of each half-alignment *A*_*L*_, *A*_*R*_ is *H* = ⌊*N/*2⌋. Positions of both half-alignments were given STR site-relative values: positions downstream to the STR site were given positive values that increase with the distance from the STR site; positions upstream to the STR were given negative values that decrease with the distance to the STR site. To analyze nucleotides forming the DNA strand on which STR takes place, nucleotides contained in *A*_*R*_ were transformed to the complementary nucleotides creating *Ã* _*R*_. Given that identical STR site-relative position exists in *A*_*L*_ and *Ã* _*R*_, nucleotide combination C is formed by a pair of nucleotides at identical positions h of the same sequence *x* (represented by a row in an alignment). *PPM*_*STR*_ is then defined as a matrix of size 16 ×*H* and is formed by nucleotide combination probabilities at position h: *PPM*_*ST R*_ = (*p*_*h*_(*c*)), *c* ∈ *C, h* ∈ *H*. The positional matrices *PPM*_*STR*_ were calculated also for the alignments of randomly selected genomic sequences. Here we refer to the probabilities derived from random matrices *PPM*_*STR*_ as *p*_*h*_(*c*|0). KLID values for *PPM*_*ST R*_ were calculated as described in Eq (11). Observed probability *p*_*n*_(*ξ*|*m*) was substituted by nucleotide combination probability *p*_*h*_(*c*) and background probability was substituted by random nucleotide combination probability *p*_*h*_(*c*|0). Results were displayed as a combined plot where bars represent the positional KLID value (*KLID*_*h*_) and colored points represent the contribution of each of the nucleotide combinations c to the *KLID*_*h*_.

### Targeting of repetitive elements

RepeatMasker annotations were obtained from UCSC goldepath as rmsk.txt tables. BED files were created from the rmsk tables using a custom awk script. BEDtools intersect tool was used to identify overlaps between IS ranges of length 26 or 27 bp and repeat annotations.

### Mapping sequences to Alu consensus

Alu repeat consensus was obtained from the publication ^50^. Bowtie2 was used to map genomic pre-integration site sequences of length 27 bp to the Alu consensus sequence. First, Bowtie2 indexes of the Alu consensus sequence were created with the bowtie-build command. Next, two rounds of mapping and filtering were run in order to select well-mapped IS sequences. In the first round, sequences were mapped to consensus with bowtie2 –f -L 5 -N 1 -i S,1,0.2 –score-min L,0,-2 –all command. Only reads mapped to a single site in consensus were selected. In the second round, bowtie2 -f -L 5 -N 1 -i S,1,0.2 –all command was run on multi mappers from the first round and alignments with a single entry were again selected. Selected alignments from both rounds were joined and converted to ranges stored in BED files. Finally, each of the ranges was transformed to a single position representing the center of the range.

### Distance to the intra-Alu sequence motif

To locate coordinates of sequence motif, seqkit locate -i -r -p CT..G…C..AG command from SeqKit tool ^59^ was used. BEDtools intersect tool was used to locate motifs positioned inside the Alu elements. To calculate distance to nearest motif to each IS, IS ranges were reduced to center position and distances were calculated using BEDtools closest tool with -D option to discriminate upstream and downstream integrations.

## Code availability

The code is available as a GitHub repository at https://github.com/dalibormiklik/IS_Motifs.git.

## Data availability

All data supporting the study can be accessed through the Open Science Framework: https://osf.io/m9hjy/?view_only=5a89ce787ce8422eaaee0c4b8fa619e4.

## Acknowledgments

We would like to thank Karen Beemon, Shelby Winans, Rik Gijsbers, and Jonas Demeulemeester for sharing the integration site data sets. We would also like to thank the members of the Laboratory of Viral and Cellular Genetics, especially Tomas Hron, Krystof Stafl, Katerina Trejbalova, and Filip Senigl for the critical reading and the discussion concerning the manuscript.

## Author contributions

D.M, J.G, and D.E conceived the study. D.M. collected and processed integration site data sets, processed EM algorithm output data, created result visualizations, and completed the manuscript. J.G. optimized and ran the EM algorithm and wrote the statistical part of the Methods. D.M., D.E., and J.H. discussed the result interpretations. All authors contributed to the final version of the manuscript.

## Extended Data

**Extended Data Table 1.** List of the data sets used in the study and sources. The bold text marks the data used to present the mixture model components in the main text. Data where the source is stated as “upon request” were requested and gained directly from the study’s authors. * number represents the sum of all IS of all variants analyzed in the present study. Numbers for individual variants can be found in Extended Data Table 2.

| Virus | Variant | Cells | genome | #IS | Publication | Source |
| --- | --- | --- | --- | --- | --- | --- |
| **HTLV-1** | **wt** | **Jurkat** | **hg19** | **4521** | **Kirk et al. 2016** | **publication** |
| **HIV-1** | **wt** | **Jurkat** | **hg38** | **161470** | **Zhyvoloup et al. 2017** | **publication** |
| **MLV** | **wt** | **Human CD34+** | **hg38** | **57991** | **De Ravin et al. 2014** | **SRR1145702** |
| **MVV** | **wt** | **HEK293T** | **hg38** | **411721** | **Ballandras-Colas et al 2022** | **GSE196040** |
| **PFV** | **wt** | **HT1080** | **hg38** | **55384** | **Lesbats et al. 2017** | **GSM2584469** |
| **ASLV** | **RCASC** | **CEF** | **galGal4** | **2096** | **Malhotra et al. 2017** | **upon request** |
| **HIV-1** | N74D | Jurkat | hg38 | 63156 | Zhyvoloup et al. 2017 | publication |
|  | wt | SupT1 | hg19 | 16803 | Vansant et al. 2020 | GSE135295 |
|  | wt (LEDGIN+) |  |  | 15683 |  |  |
|  | wt (LEDGF-KD) |  |  | 9069 |  |  |
|  | wt | SupT1 | hg18 | 6530 | Demeulemeester et al. 2014 | upon request |
|  | IN variants | SupT1/HelaP4 | hg18 | 27502 * |  |  |
| **PFV** | **wt** | **in vitro** | **hg38** | **4226638** | **Lesbats et al. 2017** | **GSM2584468** |
| **ASLV** | **ALVJ** | **CEF** | **galGal4** | 1956 | **Malhotra et al. 2017** | **upon request** |
| **ASLV** | **LR9** |  |  | 1210 |  |  |
| **ASLV** | **RCASC** | **HeLa** | **hg19** | **837** |  |  |
| **ASLV** | **RSV** | **Human CD34+** | **galGal4** | **1239** | **Moiani et al. 2014** | **SRR1282019** |

**Extended Data Table 2.** Numbers of HIV-1 total IS and IS overlapping with Alu elements. The last column displays the number of IS found inside Alu and inside the palindromic motif. Intra Alu and CT..G… C..AG columns contain two values where the first shows IS count, while the second shows the number of shuffled control IS. NA - frequencies were not calculated if the sample count was zero.

| Study | Variant | IS count | intra-Alu<br>IS; Shuffle | CT..G...C..AG<br>IS;Shuffle |
| --- | --- | --- | --- | --- |
| **Zhyvoloup** | **WT** | **161533** | **20690;16927** | **930;26.5** |
| Zhyvoloup | N74D | 63156 | 3108;6568 | 153;5 |
| **Vansant** | **WT** | **16803** | **2262;1855** | **109;1** |
| Vansant | LEDGIN | 15683 | 1652;1664 | 87;3 |
| Vansant | LEDGF-KD | 9069 | 952;897 | 44;1 |
| **Demeulemeester** | **WT** | **6530** | **697;652** | **33;0.5** |
| Demeulemeester | S119A | 1549 | 144;156 | 1;0.5 |
| Demeulemeester | S119T | 2066 | 194;208 | 2;0 |
| Demeulemeester | S119T_R231G | 2467 | 140;249 | 7;0 |
| Demeulemeester | R231G | 4897 | 357;531 | 14;1 |
| Demeulemeester | R231K | 1098 | 110;125 | 10;0 |
| Demeulemeester | R231L | 2314 | 184;239 | 3;0 |
| Demeulemeester | R231Q | 2438 | 216;273 | 6;0.5 |
| Demeulemeester | S119P | 1369 | 143;144 | 0;NA |
| Demeulemeester | S119G | 1225 | 172;117 | 0;NA |
| Demeulemeester | S119I | 1092 | 132;100 | 1;0 |
| Demeulemeester | S119K | 4682 | 436;469 | 9;1 |
| Demeulemeester | S119R | 1132 | 119;116 | 0;NA |
| Demeulemeester | S119V | 1173 | 107;112 | 0;NA |

**Extended Data Figure 1.**
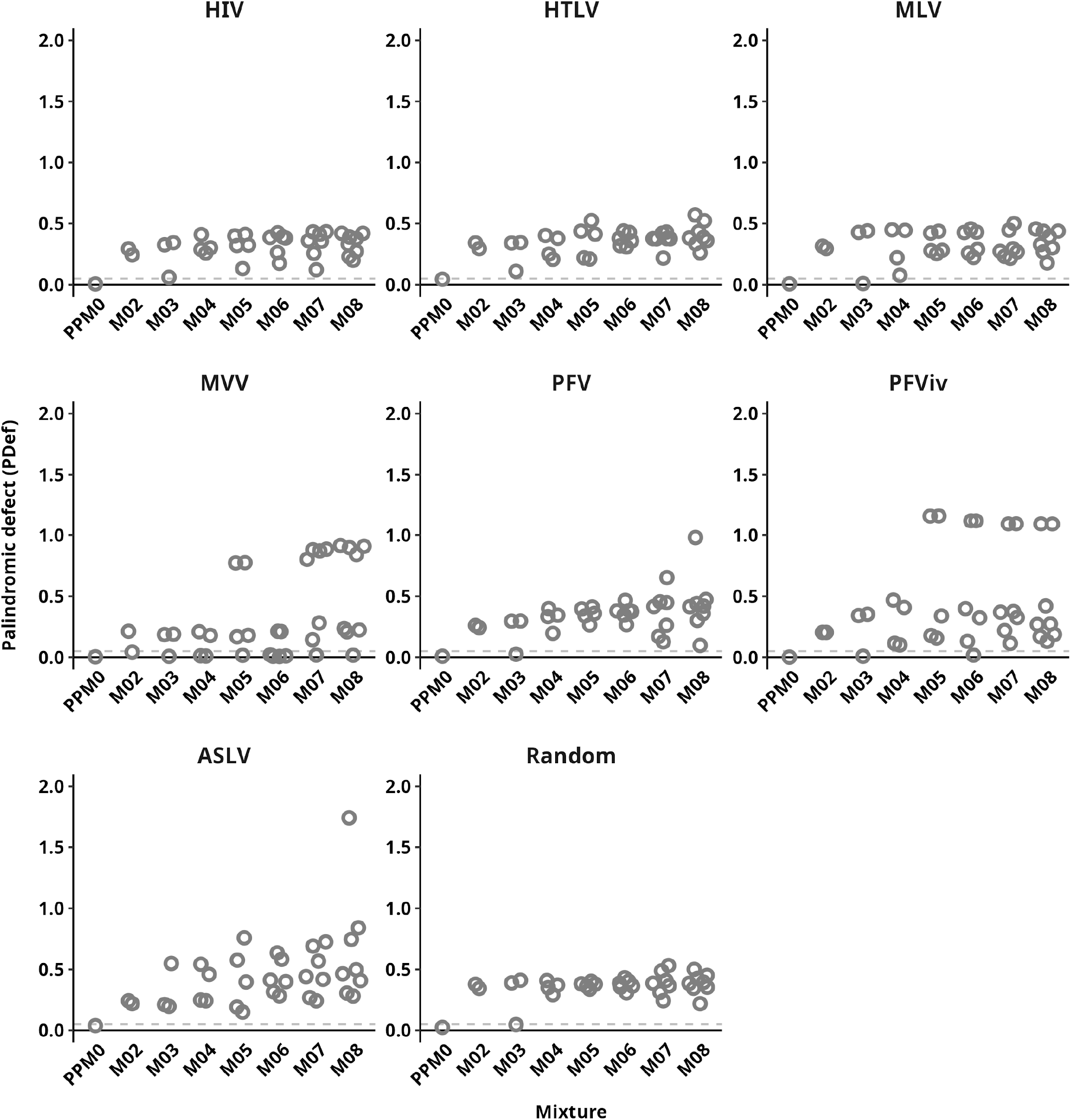
Palindromic defect (PDef) of mixture components. PDef for PPM_0_s and all components of estimated mixtures are depicted as circles. The horizontal dashed line marks the value 0.05.

**Extended Data Figure 2.**
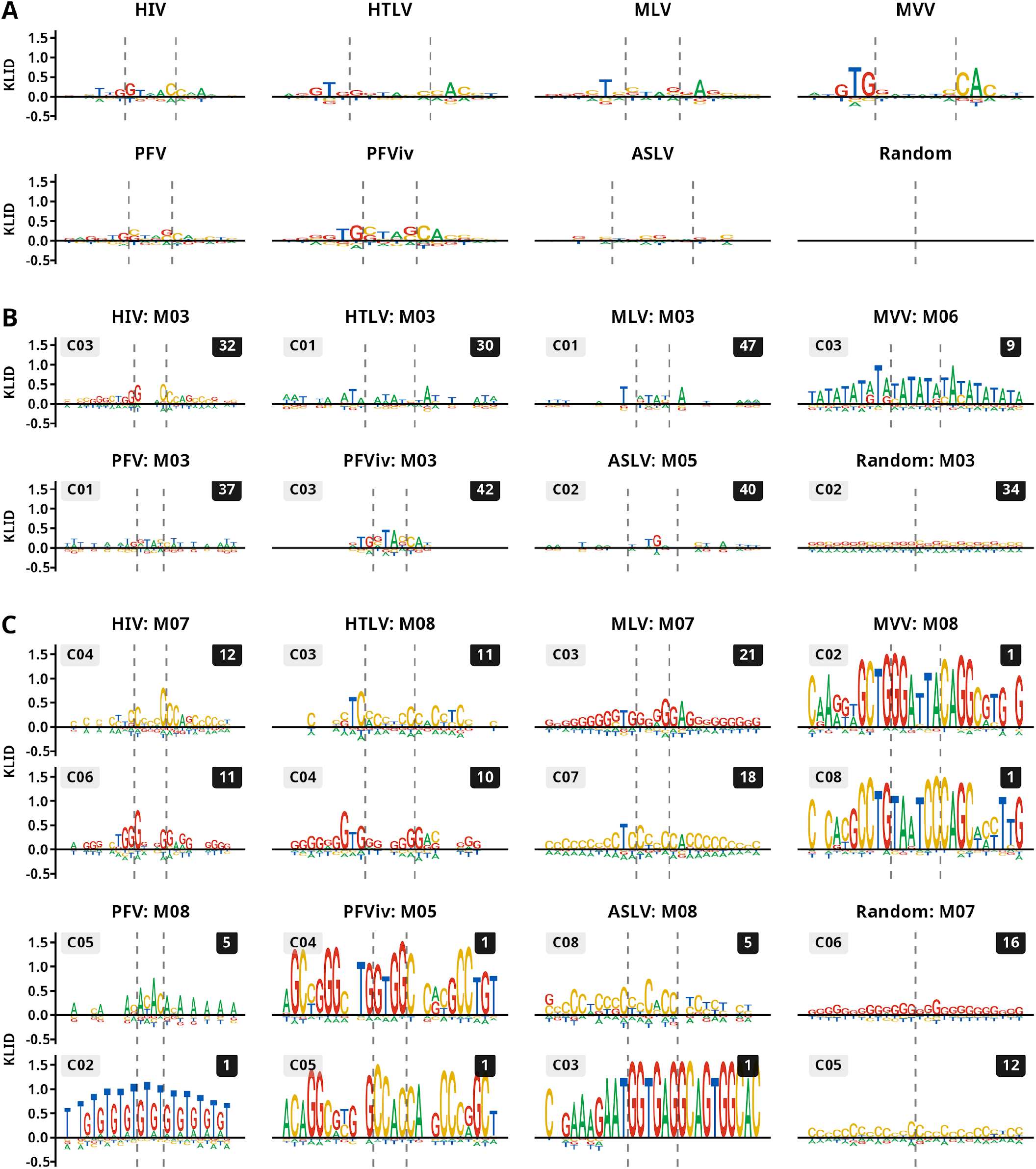
Sequence logos of selected mixture components. **A)** Sequence logos derived from PPMs representing complete sets of retroviral IS and set of random genomic sequences. Panels **B)** and **C)** show sequence logos representing **B)** component PPMs with the lowest observed PDef across all retroviral IS mixtures (i.e. the most palindromic components) and **C)** component PPMs with the highest observed PDef across all retroviral IS mixtures (i.e. the most reverse-complement asymmetric components; on top) together with the PPMs displaying the lowest reverse-complementary distance to the asymmetric PPMs. Each sequence logo contains the name of the mixture on the top, the component name on the top left corner, and the component weight multiplied by 100 on the top right corner. Logos represent IS sequences spanning 13 nucleotides to each side from the center of the sequences. Horizontal dashed lines mark sites where strand transfer reaction takes place.

**Extended Data Figure 3.**
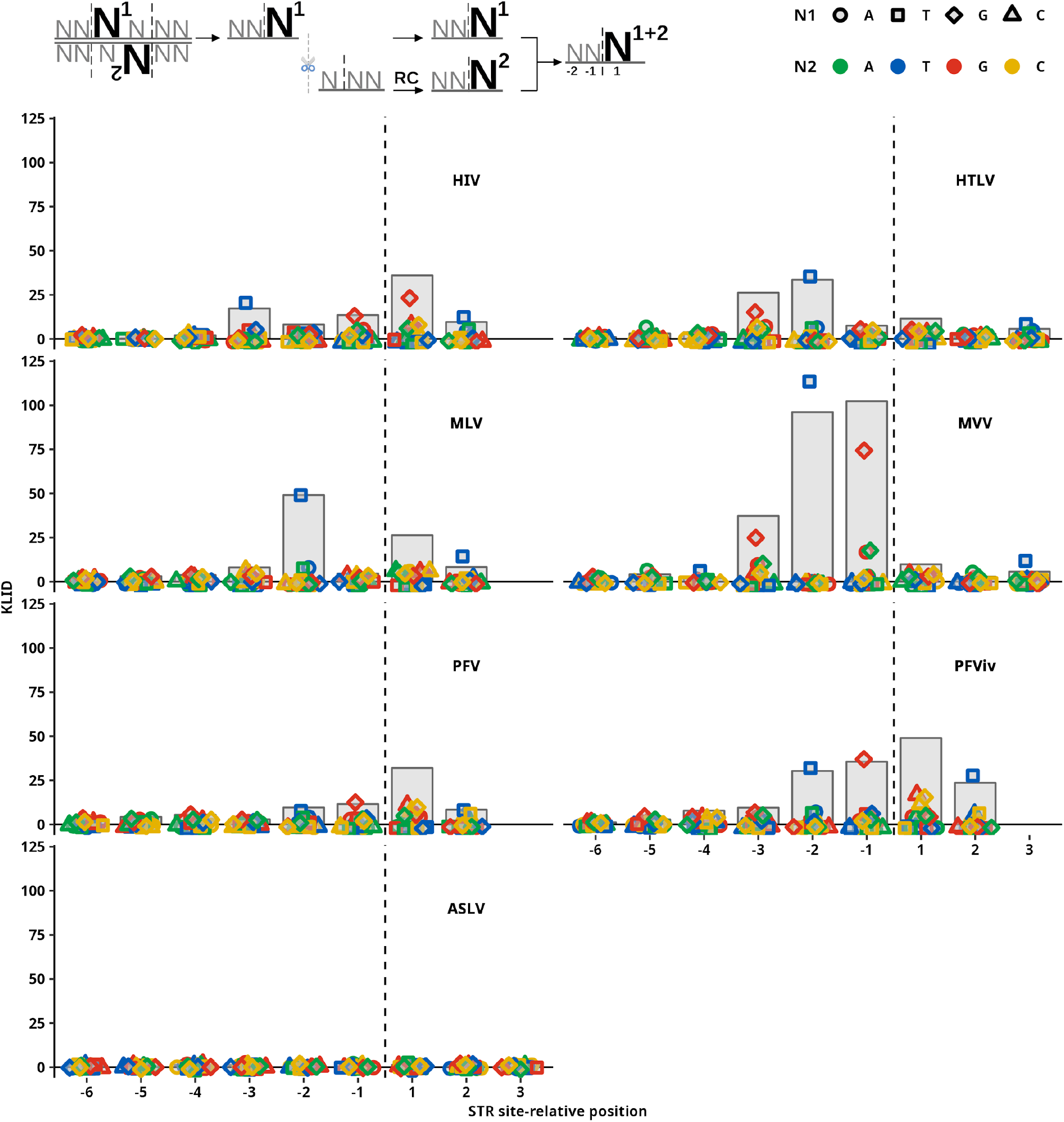
KLID representation of enrichment for positional nucleotide combinations. A scheme describing the generation of positional combination frequency at the level of a single sequence is depicted in the top left corner (RC - reverse-complement). KLID score of dinucleotide combinations at complementary sites marked by position related to the STR site. The position of the STR site is marked by the horizontal dashed line. Positions upstream to the STR site are marked with negative values. Gray bars represent the total KLID value at the position. Colored points represent individual contributions of each of the nucleotide combinations.

**Extended Data Figure 4.**
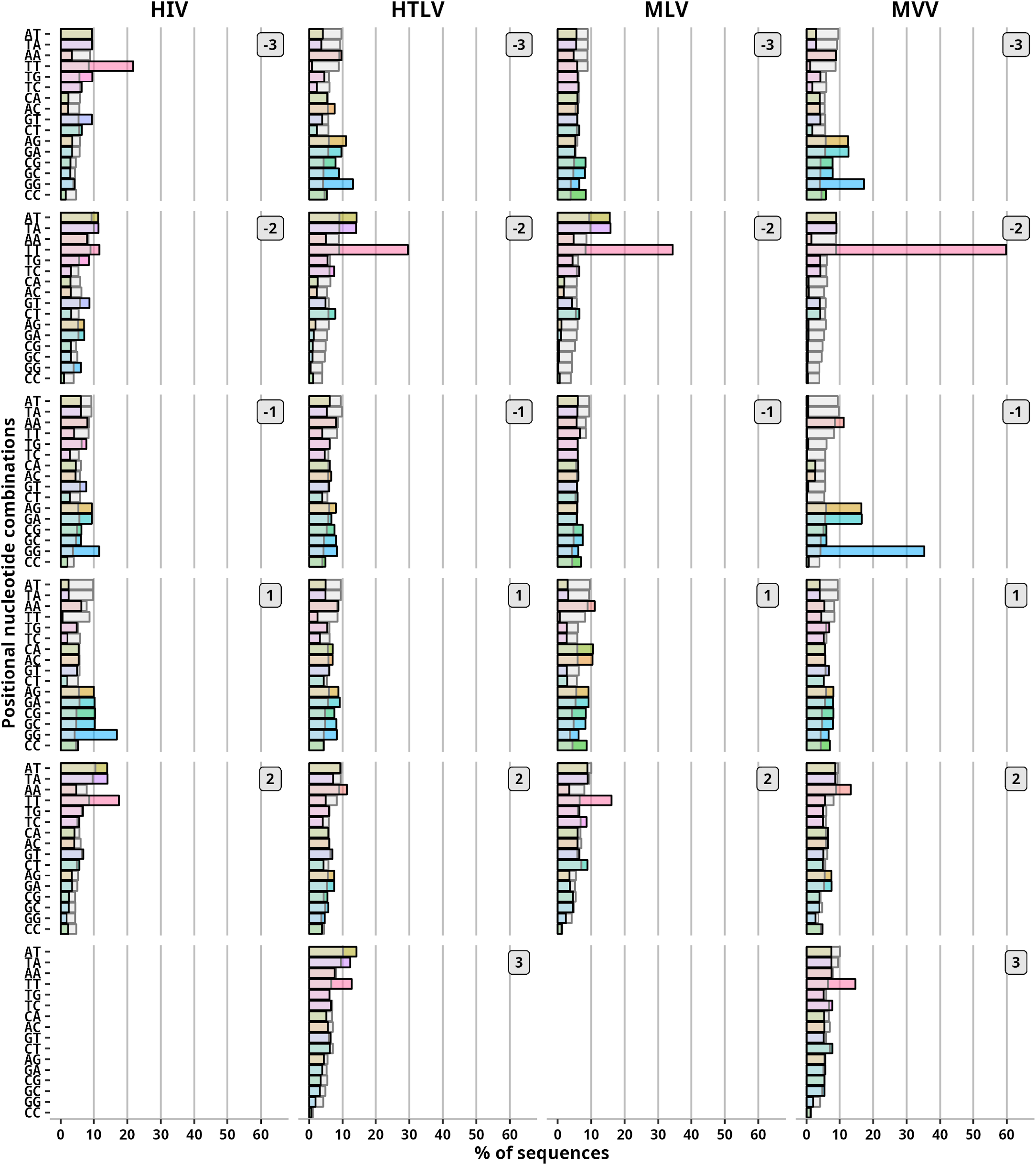
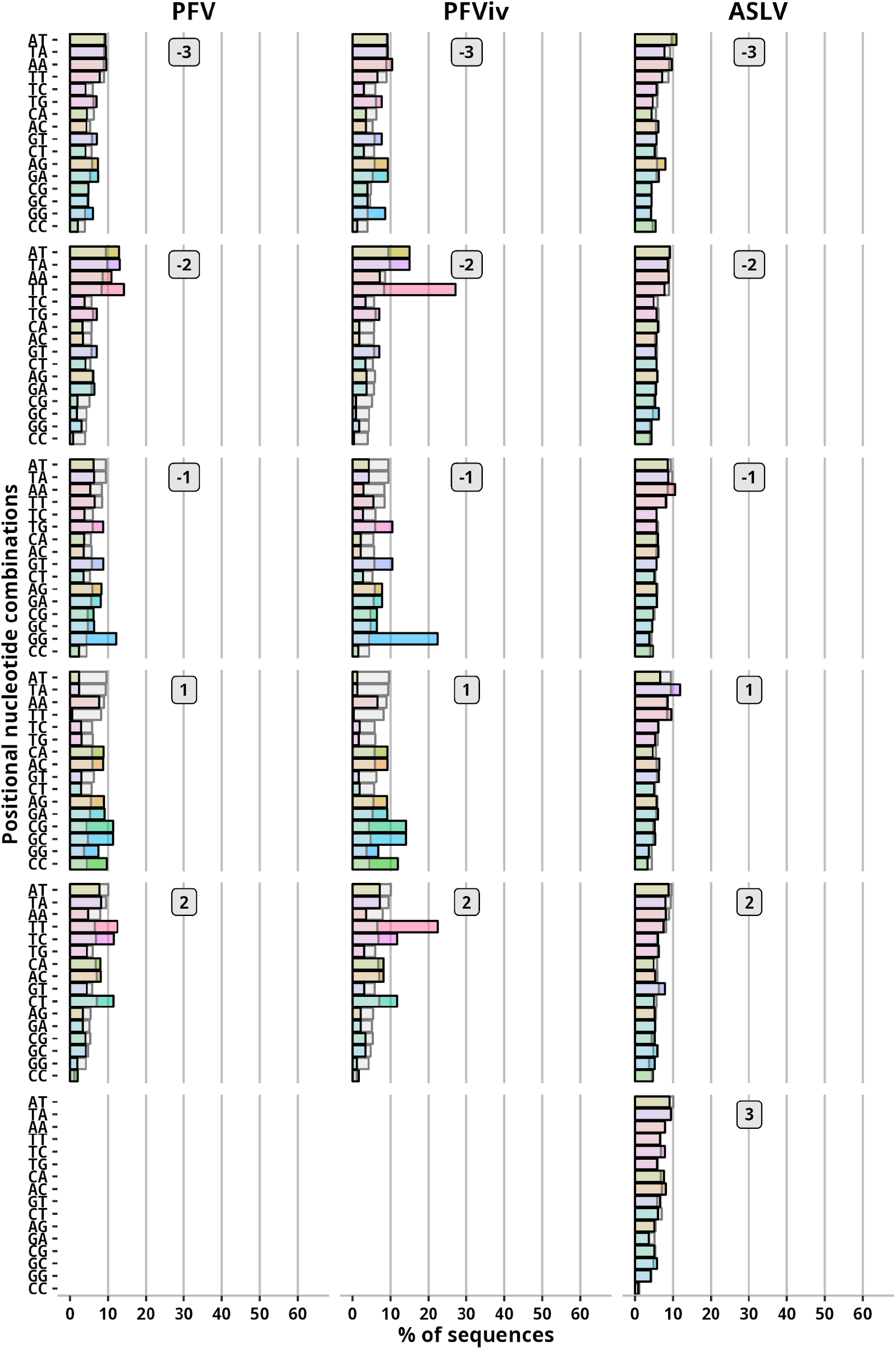
Frequency of positional combinations. Frequency of sequences with marked positional nucleotide combination. Gray bars represent frequency observed in a control (random) set of sequences. The numbers right of the bars show position relative to the cleavage site. Positions upstream to the STR site are marked with negative values.

**Extended Data Figure 5.**
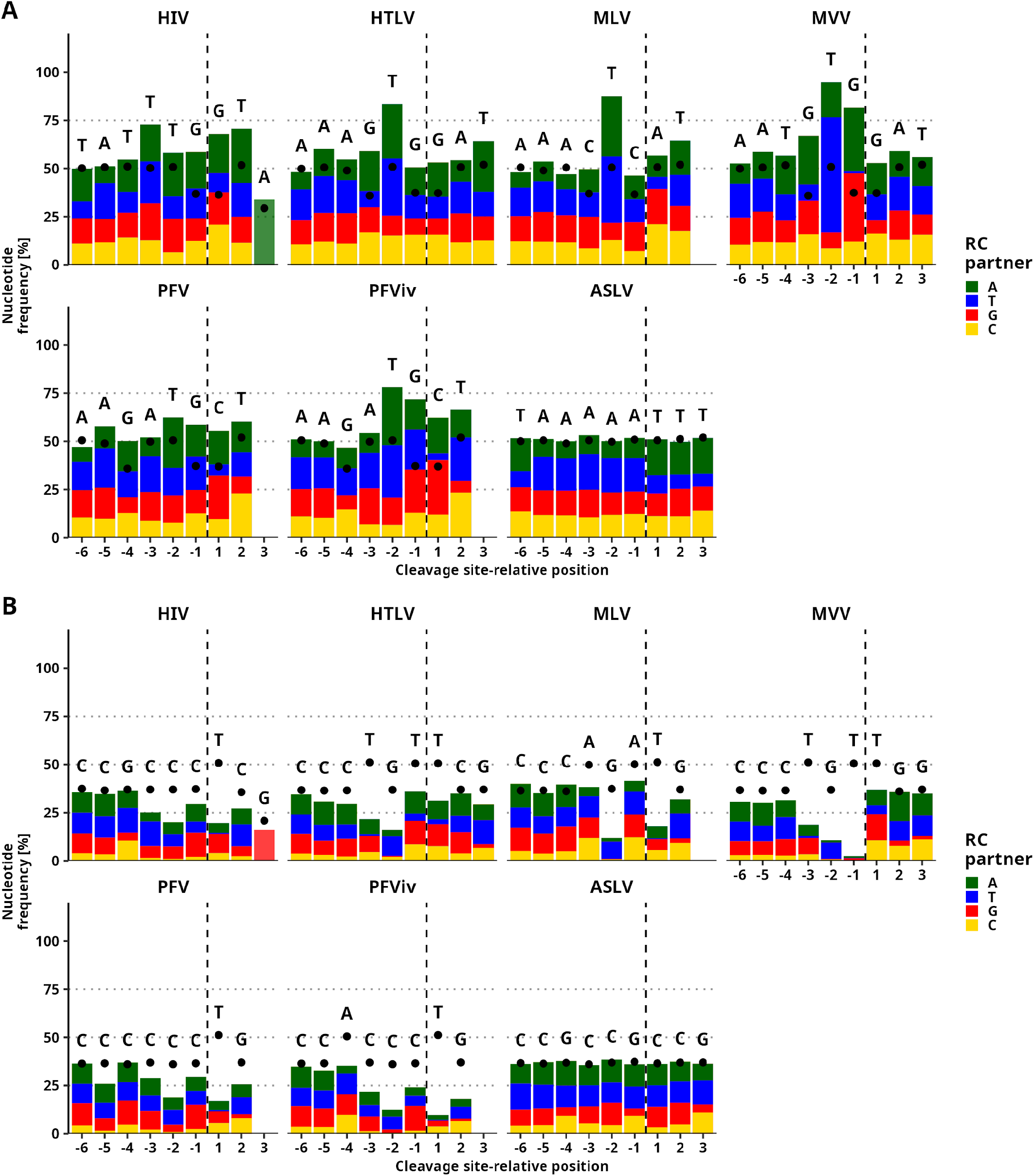
The most and least frequent nucleotides at STR site-relative positions. **A)** The most frequent nucleotide. **B)** Least frequent nucleotides. Dots represent expected frequencies at the position. The height of colored bars represents the frequency of nucleotide present at the complementary site.

**Extended Data Figure 6.**
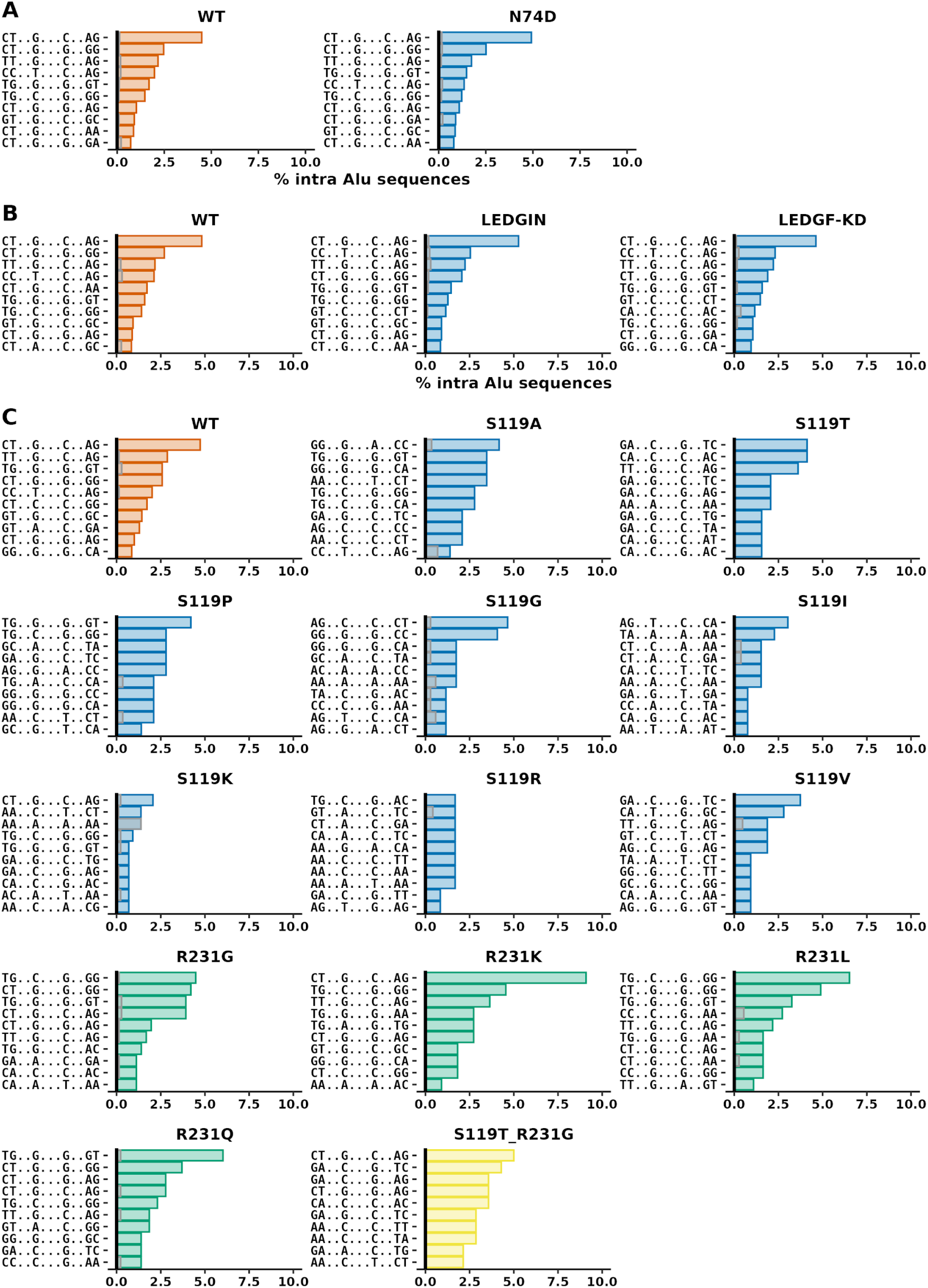
Frequency of motifs at intra-Alu IS of HIV integrations sites. Frequency of the sequence motifs among intra-Alu IS. Gray bars represent the mean frequency of 100 iterations of shuffled controls created for each IS set. The ten most frequent sequence motifs are shown. Panels **A)** and **B)** show data from IS sets where integration retargeting was achieved by interruption of capsid-CPSF6 **A)** or IN-LEDGEF/p75 **B)** interactions. Panel **C)** shows data from IS sets where mutations of S119 of R231 were introduced to HIV-1 IN. Each set contains its own set of control (wt) with unaffected integration.

**Extended Data Figure 7.**
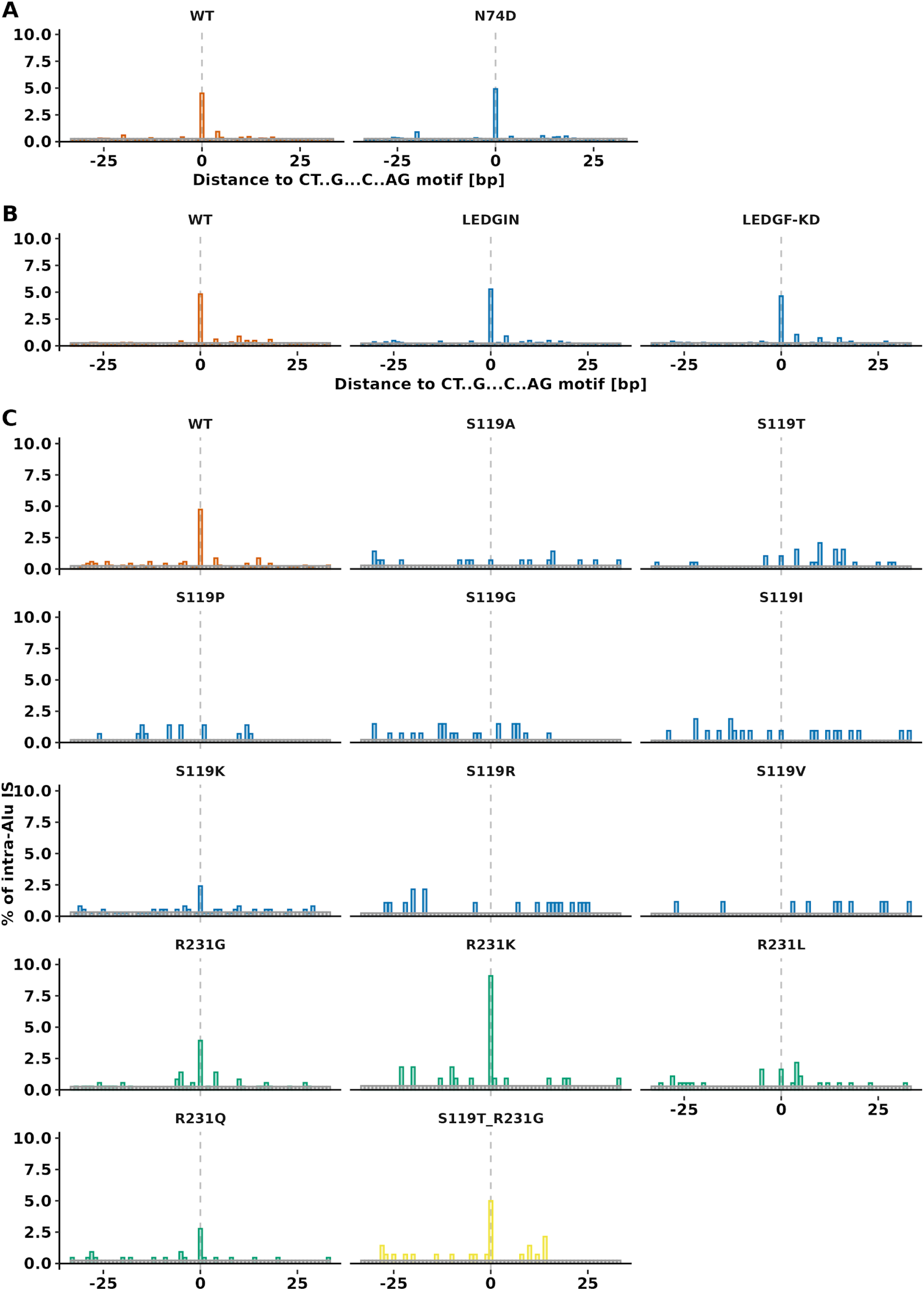
Frequency of HIV intra-Alu IS in palindromic CT..G…C..AG motif. Frequency intra-Alu IS inside and in close proximity to CT..G…C..AG palindromic motif. Gray bars represent the mean frequency of 100 iterations of shuffled controls created for each IS set. Bar plots show the frequency of intra-Alu IS 33 bp down- and upstream to the motif relative to Alu repeat orientation. Frequencies are binned by 1 bp. Panels **A)** and **B)** show data from IS sets where integration retargeting was achieved by interruption of capsid-CPSF6 **A)** or IN-LEDGEF/p75 **B)** interactions. Panel **C)** shows data from IS sets where mutations of S119 of R231 were introduced to HIV-1 IN. Each set contains its own set of control (wt) with unaffected integration.

**Extended Data Figure 8.**
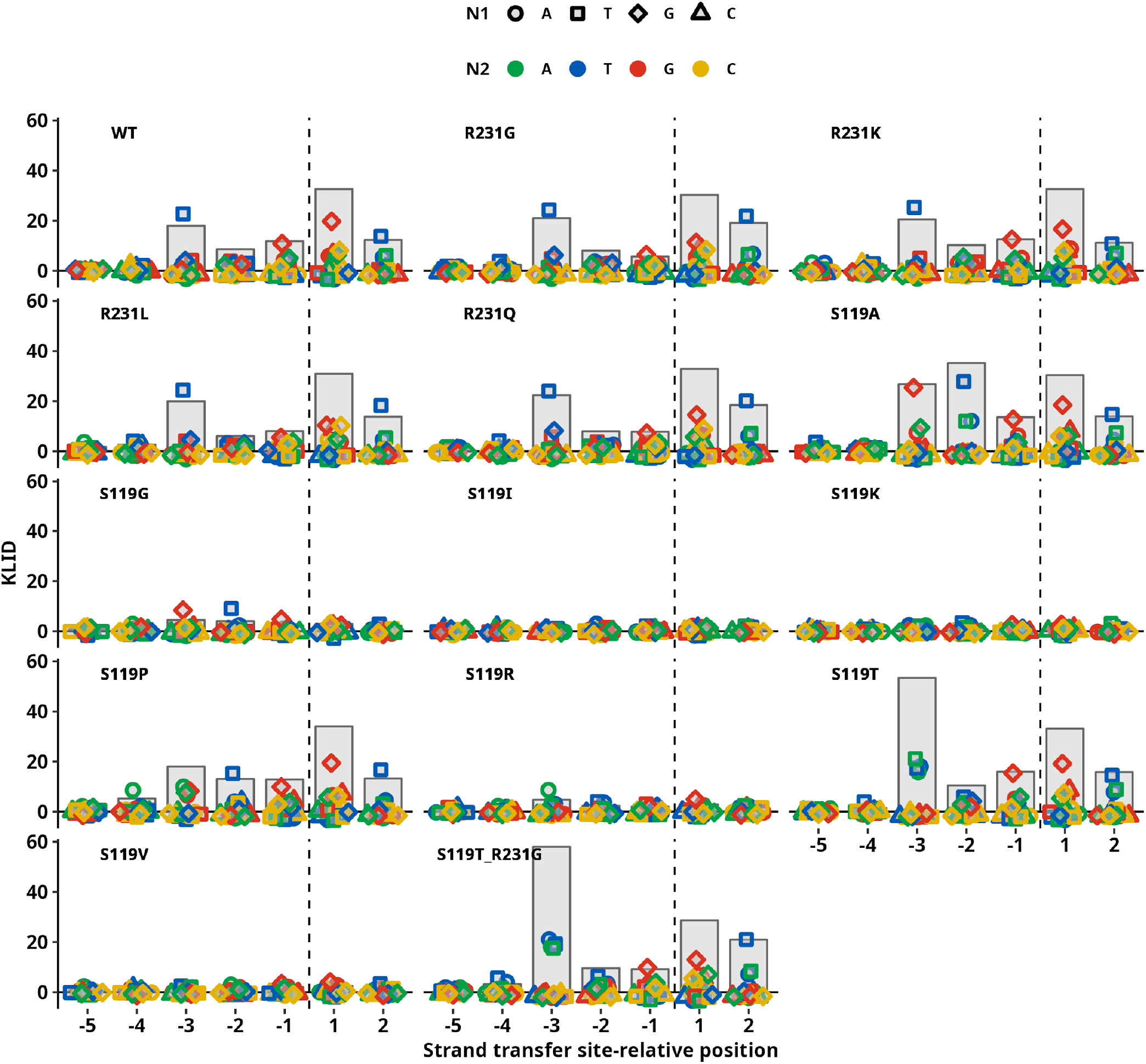
KLID representation of enrichment for positional nucleotide combinations of HIV integrase variants. KLID score of dinucleotide combinations at complementary sites marked by position related to the STR site. The position of the STR site is marked by the horizontal dashed line. Positions upstream to the STR site are marked with negative values. Gray bars represent the total KLID value at the position. Colored points represent individual contributions of each of the nucleotide combinations. Data from IS sets where mutations of S119 of R231 were introduced to HIV-1 IN.

## Supplementary Information

**Supplementary Table 1.**
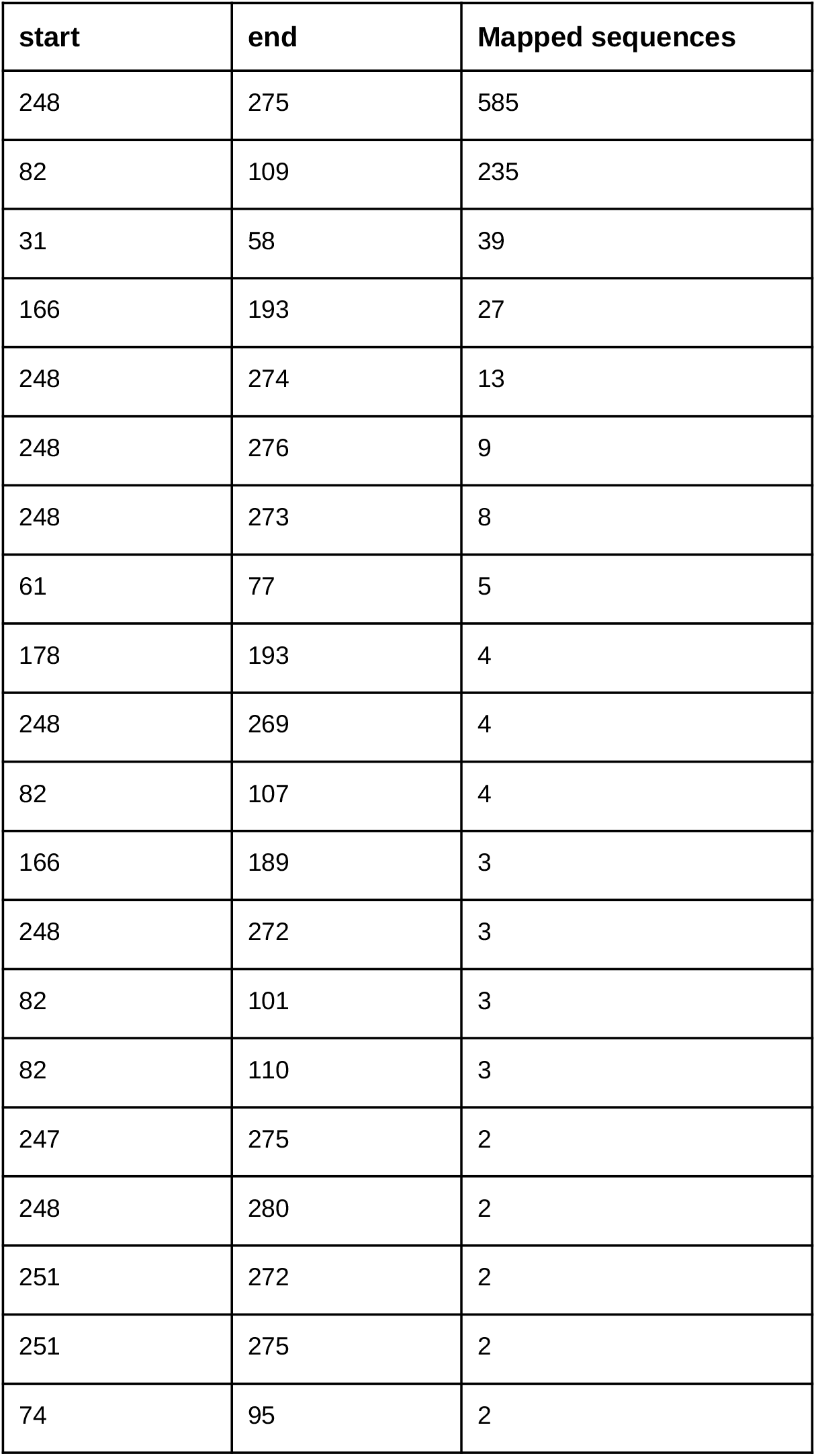
Twenty ranges in the Alu consensus sequence with the most HIV IS sequences mapped. In total, 1,587 27 bp long sequences associated with component C07 of the M08 HIV mixture were mapped to Alu consensus of which 971 sequences were identified as mapping. A full table of targeted ranges can be found in supplementary files as: *HIV_Zhyvoloup_M08_PPM07_in_hg38_rmsk_Alu_to_consensus_PosFreq*.*txt*

| start | end | Mapped sequences |
| --- | --- | --- |
| 248 | 275 | 585 |
| 82 | 109 | 235 |
| 31 | 58 | 39 |
| 166 | 193 | 27 |
| 248 | 274 | 13 |
| 248 | 276 | 9 |
| 248 | 273 | 8 |
| 61 | 77 | 5 |
| 178 | 193 | 4 |
| 248 | 269 | 4 |
| 82 | 107 | 4 |
| 166 | 189 | 3 |
| 248 | 272 | 3 |
| 82 | 101 | 3 |
| 82 | 110 | 3 |
| 247 | 275 | 2 |
| 248 | 280 | 2 |
| 251 | 272 | 2 |
| 251 | 275 | 2 |
| 74 | 95 | 2 |

**Supplementary Table 2.**
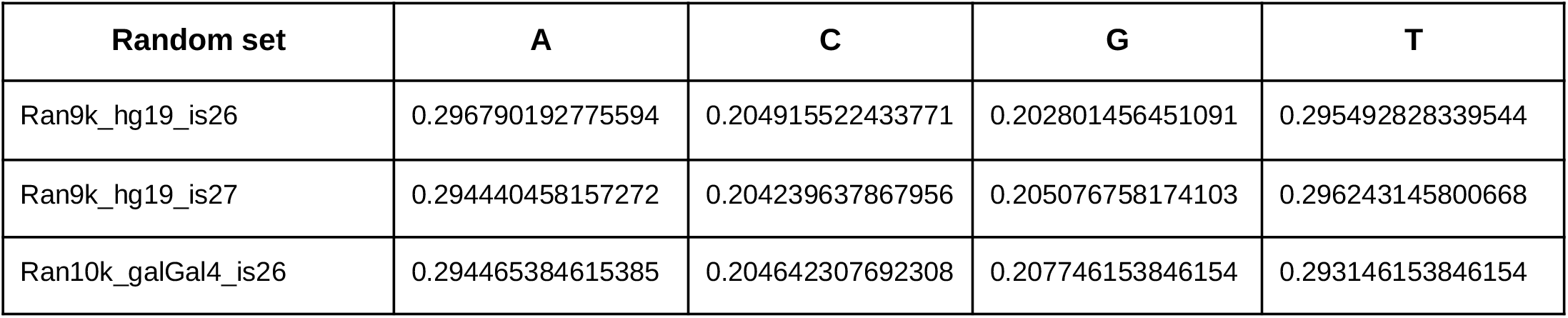
Expected nucleotide probabilities. Nucleotide frequencies were derived from 9×10^6^ (Ran9k) or 10^7^ (Ran10k) sequences of randomly selected 26 or 27 bp long ranges from human (hg19) and chicken (galGal4) reference genomes.

| Random set | A | C | G | T |
| --- | --- | --- | --- | --- |
| Ran9k_hg19_is26 | 0.296790192775594 | 0.204915522433771 | 0.202801456451091 | 0.295492828339544 |
| Ran9k_hg19_is27 | 0.294440458157272 | 0.204239637867956 | 0.205076758174103 | 0.296243145800668 |
| Ran10k_galGal4_is26 | 0.294465384615385 | 0.204642307692308 | 0.207746153846154 | 0.293146153846154 |

**Supplementary Figure 1.**
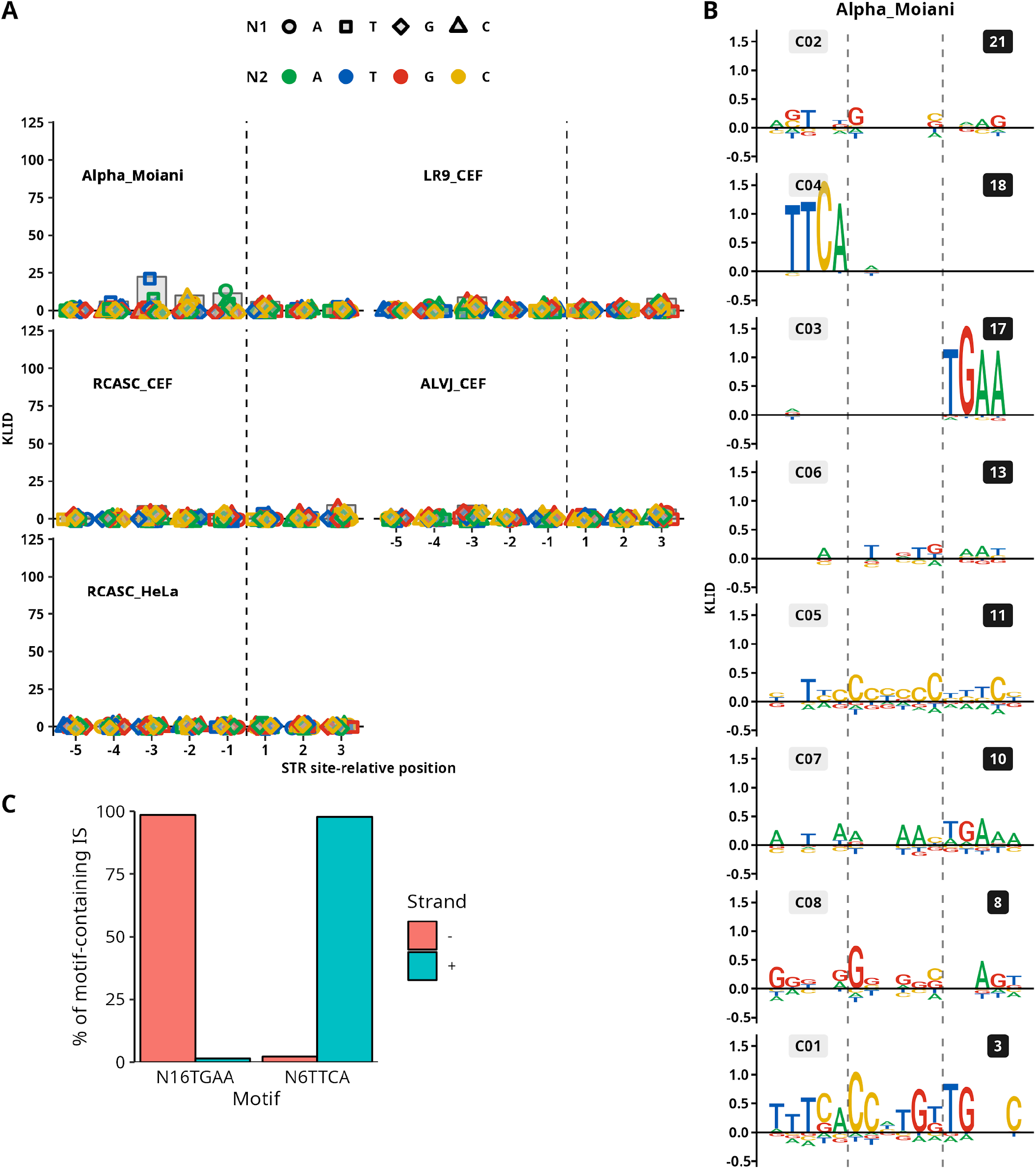
Representations of IS sequence preferences of other ASLVs. **A)** Nucleotide combinations as positional KLID values. The data set named Alpha_Moiani comes from the publication Moiani et al. 2014. Other data sets are from the work of Malhotra et al. 2017. Data sets represent distinct retroviruses or retroviral vectors transducing different cells. **B)** KLID logo representations of the M08 mixture components from the “Alpha_Moiani ‘‘ data set. Components C04 and C03 form a reverse-complementary asymmetric pair with high KLID values upstream to the STR site. **C)** Sequences containing the “N16TGAA” or “N6TTCA” motif proximal to the STR site were identified in the IS data set (number represents the number of nucleotides preceding the motif). The coordinates of the motif-containing IS were identified. Surprisingly, the majority of IS of each group motif group was mapped to either strand of the reference genome. As this finding might result from some artifact introduced during the data processing, the IS set was omitted from further analysis.

### Supplementary Methods

#### Obtaining coordinates of integration sites (IS)

Retroviral integration sites were obtained from different resources in different formats. The following text describes the procedure leading from the original file to the number-encoded IS sequence file used as input for the EM algorithm.

##### HTLV

Sequences of pre-integration genomic sequences were obtained directly from supplementary data of the publication.

##### HIV

###### Zhyvoloup et al. 2017

IS coordinates were obtained from a publication supplementary table. Replicates of DMSO-treated wt and capsid N74D mutant samples were joined to create wt and N74D IS sets. Coordinates of both IS sets were joined to create a single file.

###### Vansant et al. 2020

Coordinates were obtained from the GSE135295_ledgins1_integration_features table. Mock, LEDGIN+, and LEDGF-KD data sets were created joining bulk, GFP+, and GFP-samples from the same treatment group marked by the key present in field 8 of the feature table as follows: wt (S168, S169, S178), LEDGIN+ (S170, S171, S179), LEDGF-KD (S174, S175, S181).

###### Demeulemeester et al. 2016

Coordinates of IS in the form of text files were obtained upon request from the authors of the study. Data obtained by transduction of different cell types with identical variants were mixed. The IS coordinates of the IN variants were transformed to BED format with a custom script.

##### MLV

Raw reads were obtained from Sequence Read Archive using the fastq-dump --split-e command.

Sequences were trimmed using cutadapt with following sequences to be trimmed: LTR3nest=TGACTACCCGTCAGCGGGGGTC, LTR3rest=TTTCA,

LTR5nest=CAAACCTACAGGTGGGGTCTTTC, LTR5rest=A,

Adaptor=AGTCCCTTAAGCGGAGCCCT.

First 5’/3’ LTR sequences from primer (LTR*nest) were trimmed with

*cutadapt -m 11 -O 20 -e 0*.*1 -j 0 --trim-n*.

The rest of each LTR (LTR*rest) was trimmed using

*cutadapt -m X -O 20 -e 0 -j 0*,

where the X in -m option equals LTR*rest sequence length. The adapter was removed from sequences using

*cutadapt -m 11 -O 10 -e 0*.*1 -j 0*.

Finally, sequences containing inner proviral sequences were removed using

*cutadapt -m 11 -O 10 -e 0*.*1 -j 0*.

The resulting LTR5_F_full.fastq and LTR3_F_full.fastq were mapped to hg38 human genome assembly using bowtie2 with *-p 20 -q --no-unal -x hg38* options. Next, alignments of mapped reads were filtered to start at position 0 with *grep -v “MD:Z:0”*. Alignments with a single hit in the genome were then sorted with *grep -v “XS:i:”* command. The final BED file was created with samtools view, bedtools bamtobed, and sort commands.

##### MVV

BED formatted IS coordinates retrieved from MVV vector infected HEK293T cells were downloaded from Gene Expression Omnibus (study accession GSE196042).

##### PFV

BED formatted IS coordinates were downloaded from Gene Expression Omnibus (study accession GSE97973).

##### ASLV

###### Malhotra *et* al. 2017

SAM formatted alignments were obtained from the study’s authors upon request. First, data from experiments with identical vectors and cells were joined (48h and 120h collection time points). SAM headers from hg19 (HeLa)/galGal4 (CEF) were added to alignments for compatibility with downstream tools. Next, alignments of mapped reads were filtered to start at position 0 with *grep -v “MD:Z:0”*. Alignments with a single hit in the genome were then sorted with *grep -v “XS:i:”* command. The final BED file was created with samtools view, bedtools bamtobed, and sort commands. BED ranges were converted to contain single LTR-proximal position with custom awk script:

*awk ‘{if ($6 == “+”) print $1”\t”$1”:”$2”_”$6; else print $1”\t”$1”:”$3”_”$6;}’*

If the mapped strand was “+”, the start coordinate was used, otherwise the end coordinate was used. The awk command was followed by:

*sort* | *uniq -c* | *awk ‘{print $3”\t”$1}’* to count occurrences of each IS. Only IS coordinates with more than one occurrence were selected for further analysis.

###### Moiani *et* al. 2014

Raw reads were obtained from Sequence Read Archive using fastq-dump --split-e command. Sequences were trimmed using cutadapt with following sequences to be trimmed: LTR=TTGGTGTGCACCTGGGTTGATGGCCGGACCGTTGATTCCCTGACGACTACGAGCACCTGCATGAAGCAGAAGG

LTRend=^CTTCA

ADAPT=GTCCCTTAAC

MseADAPT=TTAGTCC

First, the majority of LTR sequence was trimmed:

*cutadapt -g $LTR -m 20 -O 30 -e 0*.*1 -j 0 --trim-n*.

Next, the end of LTR was trimmed if the sequence was exactly matching the LTRend:

*cutadapt -g $LTRend -m 15 -O 5 -e 0 -j 0*.

Finally, the adaptor sequence was removed from the reads:

*cutadapt -a $MseADAPT -m 15 -O 7 -e 0*.*1 -j 0*.

In adapter-containing sequences, the end of the read was modified to contain MseI site instead of MseI-Adapter junction using sed:

*sed ‘s/TTAGTCC$/TTAA/’*.

The reads were mapped to hg38 genome assembly using bowtie2 with *-p 20* parameter set. In the next step, alignments were filtered using custom code. First, the header of the SAM file was removed with *grep -v “^@”* and only alignments starting at position 0 were selected using *grep -v “MD:Z:0”* and only reads with a single alignment were further selected with *grep -v “XS:i:”*. The final BED file was created with SAMtools view, BEDtools bamtobed, and sort commands.

To calculate the frequency of each IS in the IS set, first, the awk was used to convert IS into ISIDs:

*awk ‘{if ($6 == “+”) print $1”\t”$1”:”$2+1”_”$6; else print $1”\t”$1”:”$3”_”$6;}’*

and then counted with:

*sort* | *uniq -c* | *awk ‘{print $3”\t”$1}’*.

IS with at least 5 occurrences were selected and the BED coordinates were transformed to represent the central position of the target site with awk:

*awk -v tsd=6 ‘BEGIN{half=tsd/2} {if ($4 ==“+”) print $1”\t”$2+half-1”\t”$2+half”\t”$4”\t”$5”\t”$6; else print $1”\t”$2-half”\t”$2-half+1”\t”$4”\t”$5”\t”$6}’*.

### Creating ranges for IS sequences

BED-formatted files containing coordinates of LTR proximal nucleotides were transformed into BED-formatted ranges covering 13 bp to each side from the center of IS (center of the inter-STR area). For this purpose, a custom awk script was used. Owing to the different origin of each IS set, the custom codes may differ and thus individual codes for each IS set is presented. Generally, *tsd* is the length of the target site duplication for a given retrovirus (or floored half of the length for HIV and MLV IS), *dist* is the final length of the sequence and *xtr* is 0 if the length of the target site duplication is even and 1 if the length of the target site duplication is odd. ISFILE is an input file with IS coordinates and output is saved into *OUTFILE*.*bed* file and sorted using *sort -k 1,1 -k2,2n ${OUTFILE}*.*bed* for compatibility with downstream tools.

#### HIV

##### Zhyvoloup et al. 2017

INVAR is either wt or n74d. This variable selects the virus variant.

*awk -v tsd=2 -v dist=26 -v xtr=1 -v invar=$INVAR ‘BEGIN{start=(dist/2)+xtr-tsd; end=tsd+ (dist/2)} $5 == invar && $3 == “+” {print $1”\t”$2-start”\t”$2+end”\t”$4”\t”“1”“\t”$4}’ ${ISFILE} > ${OUTFILE}*.*bed*

*awk -v tsd=2 -v dist=26 -v xtr=1 -v invar=$INVAR ‘BEGIN{start=(dist/2)+xtr+tsd; end=(dist/2)-tsd} $5 == invar && $3 == “-” {print $1”\t”$2-start”\t”$2+end”\t”$4”\t”“2”“\t”$4}’ ${ISFILE} >> $ {OUTFILE}*.*bed*

##### Vansant et al. 2020

*awk -v tsd=2 -v dist=26 -v xtr=1 ‘BEGIN{start=(dist/2)-xtr-tsd+1+1; end=(dist/2)+xtr+tsd-1} $6 == “-” {print $1”\t”$2-start”\t”$2+end”\t”$4”\t”$5”\t”$6}’ ${ISFILE} > ${OUTFILE}*.*bed*

*awk -v tsd=2 -v dist=26 -v xtr=1 ‘BEGIN{start=(dist/2)+xtr+tsd+1; end=(dist/2)-tsd-xtr} $6 == “+” {print $1”\t”$2-start”\t”$2+end”\t”$4”\t”$5”\t”$6}’ ${ISFILE} >> ${OUTFILE}*.*bed*

##### Demeulemeester et al. 2016

*awk -v tsd=2 -v dist=26 -v xtr=1 ‘BEGIN{start=(dist/2)+xtr-tsd; end=tsd+(dist/2)} $6 == “+” {print $1”\t”$2-start+1”\t”$2+end+1”\t”$4”\t”“1”“\t”$4}’ ${ISFILE} > ${OUTFILE}*.*bed*

*awk -v tsd=2 -v dist=26 -v xtr=1 ‘BEGIN{start=(dist/2)+xtr+tsd; end=(dist/2)-tsd} $6 == “-” {print $1”\t”$2-start”\t”$2+end”\t”$4”\t”“2”“\t”$4}’ ${ISFILE} >> ${OUTFILE}*.*bed*

#### MLV

*awk -v tsd=2 -v dist=26 -v xtr=0 ‘BEGIN{start=(dist/2)+xtr-tsd; end=(dist/2)-tsd} {print $1”\t”$2-start”\t”$3+end”\t”$4”\t”“1”“\t”$4}’ ${ISFILE} > ${OUTFILE}*.*bed*

#### MVV

*awk -v dist=26 ‘BEGIN{start=(dist/2)-1; end=(dist/2)+1} {print $1”\t”$2-start”\t”$2+end”\t”$4”\ t”“1”“\t”$6}’ ${ISFILE} > ${OUTFILE}*.*bed*

#### PFV

*awk -v tsd=4 -v dist=26 -v xtr=0 ‘BEGIN{start=(dist/2)+xtr; end=(dist/2)} {print $1”\t”$2-start”\ t”$3+end-1”\t”$5”\t”“1”“\t”$4}’ ${ISFILE} > ${OUTFILE}*.*bed*

#### ASLV

##### Malhotra *et* al. 2017

*awk -v tsd=6 -v dist=26 -v xtr=0 ‘BEGIN{start=(dist/2)+xtr-tsd; end=tsd+(dist/2)} $4 == “+” {print $1”\t”$2-start”\t”$2+end”\t”$5”\t”“1”“\t”$5}’ ${ISFILE} > ${OUTFILE}*.*bed*

*awk -v tsd=6 -v dist=26 -v xtr=0 ‘BEGIN{start=(dist/2)+xtr+tsd; end=(dist/2)-tsd} $4 == “-” {print $1”\t”$2-start”\t”$2+end”\t”$5”\t”“2”“\t”$5}’ ${ISFILE} >> ${OUTFILE}*.*bed*

##### Moiani *et* al. 2014

*awk -v tsd=6 -v dist=26 -v xtr=0 ‘BEGIN{start=(dist/2)+xtr; end=(dist/2)} {print $1”\t”$2-start”\ t”$3+end-1”\t”$5”\t”“1”“\t”$4}’ ${ISFILE} > ${OUTFILE}*.*bed*

### Obtaining genomic sequences from IS ranges

BEDtools getfasta tool was used to retrieve genomic sequences. IS BED ranges and FASTA files with sequences of chromosomes served as input. In resulting FASTA files, all nucleotides were converted to capitals with

*cat ${ISSEQ}*.*fa* | *sed ‘s/a/A/g’* | *sed ‘s/c/C/g’* | *sed ‘s/g/G/g’* | *sed ‘s/t/T/g’ > ${ISSEQ}_CAP*.*fa*,

Where ISSEQ is an output file from the getfasta tool.

The right alignment of the sequences was inspected through the classical sequence logo produced by the ggseqlogo R package.

### Converting sequences to numeric strings

To use sequences as input for the EM algorithm, nucleotide sequences needed to be converted to a string of numbers. Thus, each nucleotide was given a numeric code. Sequences in the FASTA file were first converted to a table, where sequences are placed into a single column and each sequence occupies a single row:

*grep -v “>“ ${ISSEQ}_CAP*.*fa > ${ISSEQ}_is26*.*txt*.

Finally the sequences were converted to numeric strings:

*sed ‘s/[Aa]/1/g’ ${ISSEQ}_is26*.*txt* | *sed ‘s/[Cc]/2/g’* | *sed ‘s/[Gg]/3/g’* | *sed ‘s/[Tt]/4/g’* | *sed ‘s/[Nn]/5/g’ ${ISSEQ}_is26_num*.*txt*

